# Synergistic application of continuous granulation and selective laser sintering 3D printing for the development of pharmaceutical dosage forms with enhanced dissolution rates and physical properties

**DOI:** 10.1101/2021.02.13.430988

**Authors:** Rishi Thakkar, Yu Zhang, Jiaxiang Zhang, Mohammed Maniruzzaman

## Abstract

This study demonstrated the first case of combining novel continuous granulation with powder-based pharmaceutical 3-dimensional (3D) printing processes to enhance the dissolution rate and physical properties of a poorly water-soluble drug. Powder bed fusion (PBF) and binder jetting 3D printing processes have gained much attention in pharmaceutical dosage form manufacturing in recent times. Although powder bed-based 3D printing platforms have been known to face printing and uniformity problems due to the inherent poor flow properties of the pharmaceutical physical mixtures (feedstock). Moreover, techniques such as binder jetting currently do not provide any solubility benefits to active pharmaceutical ingredients (APIs) with poor aqueous solubility (>40% of marketed drugs). For this study, a hot-melt extrusion-based versatile granulation process equipped with UV-Vis process analytical technology (PAT) tools for the in-line monitoring of critical quality attributes (i.e., solid-state) of indomethacin was developed. The collected granules with enhanced flow properties were mixed with vinylpyrrolidone-vinyl acetate copolymer and a conductive excipient for efficient sintering. These mixtures were further characterized for their bulk properties observing an excellent flow and later subjected to a PBF-3D printing process. The physical mixtures, processed granules, and printed tablets were characterized using conventional as well as advanced solid-state characterization. These characterizations revealed the amorphous nature of the drug in the processed granules and printed tablets. Further, the *in vitro* release testing of the tablets with produced granules as a reference standard depicted a notable solubility advantage (100% drug released in 5 minutes at >pH 6.8) over the pure drug and the physical mixture. Our developed system known as DosePlus combines innovative continuous granulation and PBF-3D printing process which can potentially improve the physical properties of the bulk drug and formulations in comparison to when used in isolation. This process can further find application in continuous manufacturing of granules and additive manufacturing of pharmaceuticals to produce dosage forms with excellent uniformity and solubility advantage.

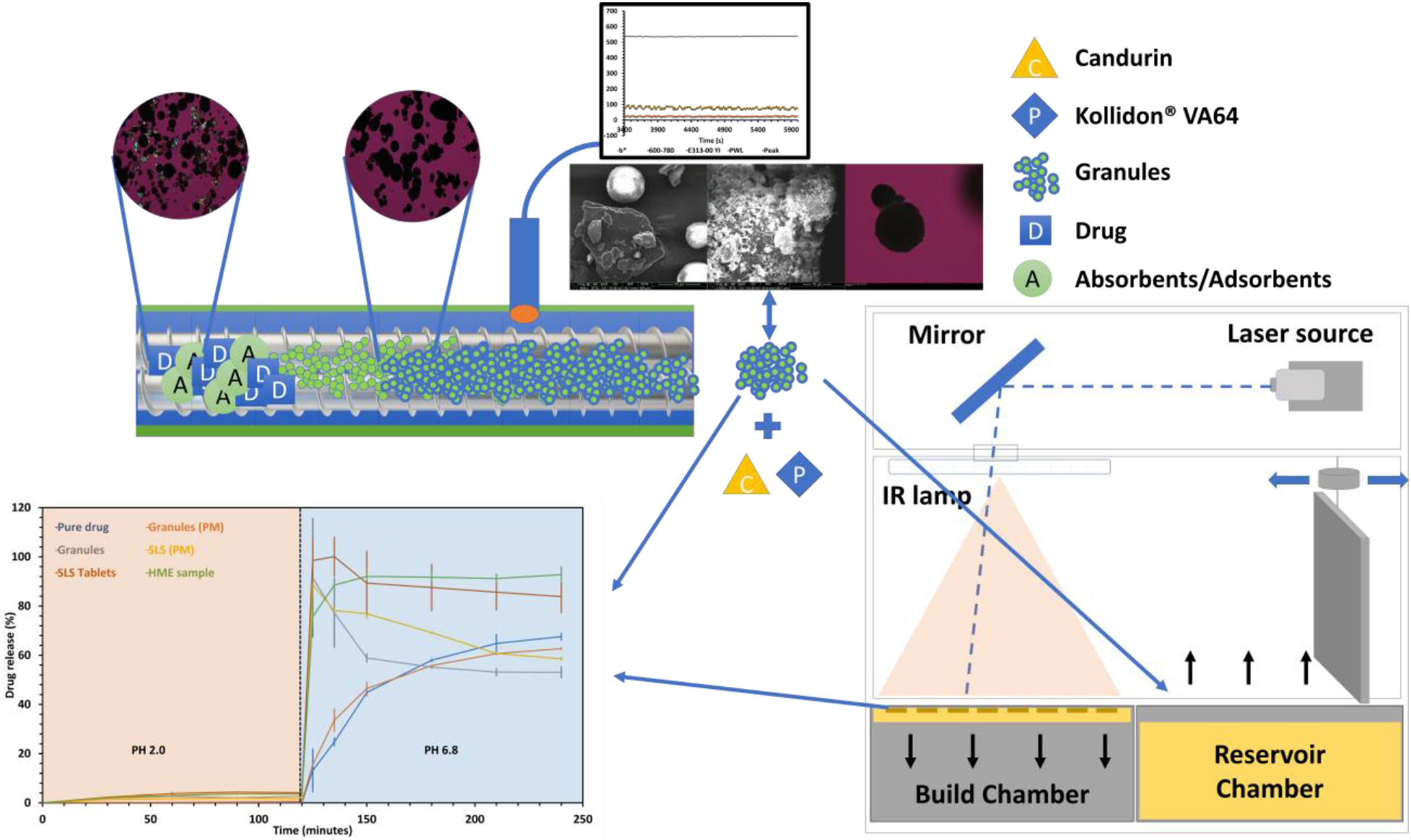

## 1. Introduction

Additive manufacturing (AM) has gained much attention amongst researchers across the field of engineering and technology since the early 1970s [1,2]. This interest has extended to the pharmaceutical field in the past two decades mainly due to the versatility of designing [3,4], rapid prototyping [5], and the possibility of 3-dimensional (3D) printing of personalized (patient-specific) dosage forms [6,7]. AM techniques use ‘STL’ (standard tessellation language) or ‘AMF’ (additive manufacturing file) formats by slicing them into g-code (x, y, z coordinates for constructing 3D computer-aided design (CAD) constructs) [8]. Pharmaceutical researchers have employed material extrusion techniques such as fused deposition modeling (FDM)/fused filament fabrication (FFF) [9], and semi-solid extrusion (SSE) [10] due to their inherent advantage of complex structure designing. On the other hand, powder bed-based 3D printing such as binder jetting, and powder bed fusion (PBF) has found application in creating highly porous dosage forms where the powder bed supports the 3D constructs [11,12]. Each of these enlisted techniques have specific material considerations where FDM utilizes thermoplastic polymeric filaments with optimum hardness and flexibility [13], SSE requires solvent considerations to prevent structure deformation post extrusion [14], binder jetting needs extensive binder formulation considerations to prevent print head overheating [2], and PBF requires material consideration to prevent drug substance degradation due to increased temperature and high power lasers [15]. Apart from individual material considerations, both powder bed-based techniques require a power batch with excellent flow properties, uniform particle size distribution, and uniform drug distribution [11].

Flow properties are crucial in the powder bed-based techniques, as feedstock from the feed region is repeatedly conveyed to the powder bed where the particles are either combined by the means of a binder (binder jetting) or are fused using a high power laser in laser sintering (LS) [16,17]. Particle size distribution holds importance as agglomerates/aggregates can create print defects and disturb the build surface leading to print failures [11]. Moreover, both these processes expose the feed region to vibrations. The difference in particle densities, shape, size of the blend components (drug substance, polymer, excipients) can lead to component segregation which can, in turn, impact the drug content of the manufactured dosage forms [18]. Furthermore, these 3D printing techniques work best with materials with particle sizes less than or equal to 100μm, for instance commercially available polyamide (PA 12) suitable for LS processes has a size range of 45-90μm [19–21].

Shear cell testers have been used to evaluate the flow properties of pharmaceutical compositions. They produce consolidated shear initially (pre-shear), and then expose the sample to a series of normal stresses in a range while recording the corresponding shear stresses (the yield locus) [22]. The yield locus analysis is used to determine the angle of internal friction and cohesion for a sample material which then calculates the overall strength of the sample under compressive load. The points observed can be imagined as the shear stress required for the flow to initiate at a given normal stress. The flow function coefficient derived (ff_c_=σ_1/_ σ_c_) is the ratio between σ_1_ (consolidation stress) and σ_c_ (unconfined yield strength) at a particular pre-consolidation condition. A smaller ff_c_ corresponds to poor flow which means the powder components are more cohesive. Megarry A. J. et al., (2019) used a historical data set (3909 experiments) from a shear cell apparatus to establish the general flow properties of pharmaceutical compositions (pure drug substances, blends, granules). Megarry A. J. and colleagues further found that out of the 199 unique pure drug substance samples, >66% samples had an ff_c_ value of <4, and >50% samples had an ff_c_ value of <2, i.e., they had extremely poor flow properties. Whereas >87% blends had ff_c_ values of >4 with most blends within the values of 4-6. The study also depicted that ff_c_ values of >10 (excellent flow) were predominantly observed for granules [22]. This case study observed that pharmaceutical drug substances generally have poor flow properties and thereby require additives, or additional processing (granulation, micronization) to facilitate the efficient mass transfer, which is essential in pharmaceutical unit processes.

In conventional pharmaceutical processes, granulation techniques have been utilized to solve these mass transfer issues [23–25]. Unfortunately, conventional pharmaceutical granulation methods (wet granulation, dry granulation, melt granulation) are not particularly suitable for the 3D printing techniques under discussion as the produced granules from such processes are 200-400μm in size [26]. With the upsurge of interest in the pharmaceutical applicability of powder bed-based techniques, research on processes to develop materials having the capability to circumvent the aforementioned mass transfer challenges is of paramount importance. This study discusses an inventive technique for manufacturing granules suitable for the discussed 3D printing processes.

Currently, the only 3D printed (binder jetting) dosage form approved by the United States food and drugs administration (USFDA) contains levetiracetam which is highly water-soluble (1.04g/mL) pyrrolidine with anti-epileptic activity under the brand name Spiritam® [27]. The technique manufactures highly porous tablets with rapid disintegration time and finds applicability for drugs with disintegration limited absorption. This means the drug gets absorbed as soon as the dosage form disintegrates as it has high water solubility and permeability [28]. These drugs fall under class I of the biopharmaceutical classification system [29]. The powder bed-based 3D printing techniques have not found any applicability in improving the solubility of BCS class II (low solubility, high permeability) and IV (low solubility, low permeability) yet. BCS class II drugs account for >40% of drug substances, these drugs suffer dissolution limited absorption i.e. their solubility is the rate-limiting step for their absorption and thereby their therapeutic action [30]. This directs the discussion to the second objective of this study where the manufactured granules demonstrate a significant increase in their dissolution rate as compared to their crystalline counterpart and unprocessed blends.

For this study indomethacin (IND) was selected as a model drug. IND is a non-steroidal anti-inflammatory drug (NSAID) that inhibits cyclooxygenase (COX-I and COX-II) enzymes non-selectively [31]. IND is deemed practically insoluble in water (0.937e^−6^g/mL), moreover, it also exhibits poor flow properties because of its highly irregular crystal habits and inter particulate cohesion [32,33]. Thereby IND was the ideal candidate to test the granulation technique’s applicability for powder bed-based 3D printing (PBF) of drug substances with poor solubility and flow properties. The drug crystals were completely fragmented into their amorphous form by the means of heat and shear-induced by twin-screw processing and absorbed onto the surface of highly porous silicates, this process was monitored using an in-line ultraviolet-visible (UV-Vis) reflectance probe. The manufactured granules were blended with a thermoplastic vinyl pyrrolidone-vinyl acetate copolymer (Kollidon® VA 64) and potassium aluminum silicate-based pearlescent pigment (Candurin®) which is a food-grade visible laser absorbing thermally conductive excipient. This blend was processed into tablets using LS 3D printing. The manufactured tablets released >80% of their drug load in <5 minutes into the dissolution testing apparatus, whereas its crystalline counterpart released ≅60% of an equivalent drug load over two hours.

## 2. Material and methods

### 2.1 Materials

Indomethacin (Tokyo Chemical Industries, Lot no. D3NIJJR), magnesium aluminometasilicate (Neusilin US 2, Lot no. 901002, Fuji chemical industries co., ltd Toyama pref., Japan), silicon dioxide (Fujisil™, Lot no. 906003, Fuji chemical industries co., ltd Toyama pref., Japan), polysorbate 80 (Lot no. BCCB4768, Sigma-Aldrich^®^, Missouri, USA), vinyl pyrrolidone-vinyl acetate copolymer (Kollidon^®^ VA 64, Lot no. 94189624U0, BASF, Ludwigshafen, Germany), potassium aluminum silicate-based pearlescent pigment (Candurin^®^, Lot no. W150645X08, Merck KGaA, Darmstadt, Germany), HPLC grade acetonitrile was purchased from Fisher Scientific (Pittsburg, PA); all other chemicals and reagents were ACS grade or higher.

### 2.2 HME based granulation process

To ensure the reproducibility of the process, three batches of the physical mixture were prepared using the geometric dilution technique. Each 200g batch of the physical mixture contained a 40% IND drug load, 27.5% of each of the inorganic highly porous absorbents (silicon dioxide and magnesium aluminometasilicate), and 5% of polysorbate 80 (non-ionic surfactant) (here on out this composition will be referred as PM-I). This mixture was transferred to a twin-screw gravimetric feeder with stirring agitators (Brabender Technologie, Ontario, Canada) which was calibrated for the blend to quantify and control the amount of feed going into the system, post-calibration the feed rate was set to 5g/min. The feed was processed using a twin-screw extruder with a 12mm outer diameter (OD) (ZSE 12 HP-PH, Leistritz Advanced Technologies Corp., Nuremberg, Germany). The temperature for each zone has been outlined in fig. 1 along with other processing parameters required to define the process. The granules were collected after the process was stabilized. The physical mixture and the collected granules were subjected to bulk property testing, a series of solid-state characterizations, and performance testing before using them for LS 3D printing.

**Figure 1.**
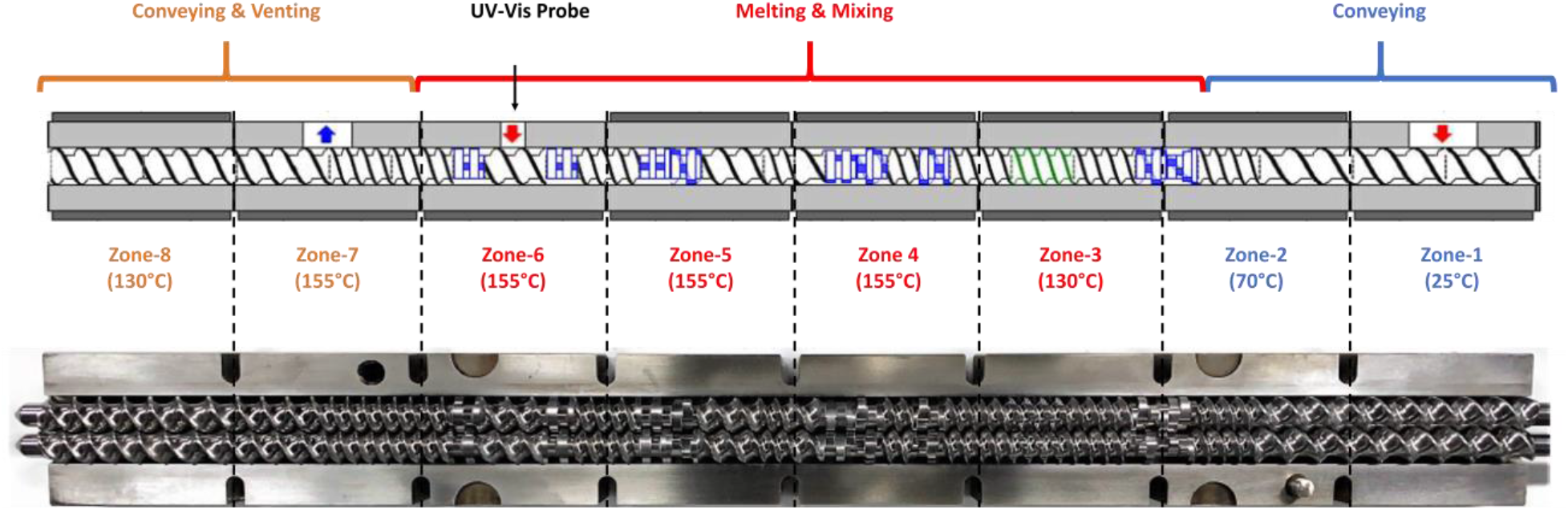
A schematic of the processing conditions employed during the granulation process.

### 2.3 In-line process monitoring

The aforementioned HME processing parameters were optimized by using a UV-vis reflectance probe with a 316L Stainless Steel/Nickel alloy tip and sapphire window. The probe was later used as a process analytical tool (PAT) for monitoring the uniformity and amorphous conversion of the subsequent batches (Equitech Int’l Corporation, New Jersey, USA). Indomethacin has a unique property, in its crystalline γ-form, it exhibits a pinkish white appearance, whereas on amorphous conversion its color changes to yellow [34]. During the granulation process, CIELAB yellow-blue color space coordinate (b*), custom selected wavelength (600-700 nm), yellowness index (‘E313-00 YI’ which is supposed to trend with b*), the wavelength of maximum reflectance over the measured region (PWL), and reflectance value at the PWL (Peak) were observed by the reflectance probe and were used as an indicator of amorphous conversion and inspect the stability of the process. The physical mixture was processed with the probe in place with different temperature conditions ranging from 140-155°C (below indomethacin’s melting point), the samples were collected, and the yellowness values attained from the probe were noted. These samples were tested using powder X-ray diffraction (pXRD) analysis. Processing conditions where the samples observed no crystalline peaks were selected and the corresponding yellowness values were used to observe the uniformity of subsequent processes.

### 2.4 Bulk properties testing

#### 2.4.1 Digital light microscopy

Digital microscopy was used to investigate the morphology of the drug crystals, PM-I, and manufactured granules (Dino light, Torrance, California, USA). The microscope was set to a magnification of 65X which was sufficient to observe the particle characteristics of samples. This technique was used as a convenient quality control tool to observe the absence of any drug crystals in the manufactured granules post-processing. It was also used to understand the crystal morphology which provided deeper insight into the flow characteristics of the drug. Although digital microscopy was suitable for investigating particle morphology, it was not suitable to study and observe the particle surface pre- and post-processing. Moreover, it did not provide any insight into the solid-state of the drug in the samples. Hence, polarized light microscopy and scanning electron microscopy were conducted for all the aforementioned samples to further investigate their surface properties.

#### 2.4.2 Polarized light microscopy (PLM)

Polarized light microscopy using an Olympus BX53 polarizing photomicroscope (Olympus America Inc., Webster, TX, USA) was used to investigate the crystallinity of IND, PM-I, and the granules. The microscope had a Bertrand Lens and a 10X objective lens. The samples were evenly dispensed on a glass slide which was later dusted off to remove excess powder and a coverslip was placed onto it. The sample slides were then observed using a 10X magnification lens and an appropriate zone was selected to observe the state of the sample. The magnification was then increased to 20X to further observe the crystals with more clarity. Crystalline particles possess the property of birefringence, which is characteristic of crystalline substances, hence it was predicted that the granules will not depict any birefringence. After focusing on the sample, snapshots were taken with a QICAM Fast 1394 digital camera (QImaging, BC, Canada). These images were taken with and without a 530 nm compensator (U-TP530, Olympus^®^ corporation, Shinjuku City, Tokyo, Japan). The snapshots were processed using Linksys 32 software^®^ (Linkam Scientific Instruments Ltd, Tadworth, UK).

#### 2.4.3 Scanning electron microscopy

To understand the surface morphology of the drug crystals, physical mixture, and the processed granules, a scanning electron microscope (Quanta FEG 650 ESEM, FEI Company, Hillsboro, OR, USA) was used. The samples were first exposed to vacuum gold sputtering (EMS Sputter Coater, Hatfield, PA, USA) before observing them under the microscope. Microscopic images were captured at an accelerated voltage of 10 KV, emission current of 15μÅ, the working distance of ≈10 mm, and a spot size of 3. The magnification was varied from 100X to 2000X based on the purpose of the observation.

#### 2.4.4 Powder flow

A United States Pharmacopoeia (USP) compliant flowability tester, with funnel attachments (BEP2, Copley Scientific Limited, Nottingham, UK) was used for the flow-through orifice study. The purpose of this test was to stimulate the flow in a hopper or other mass transfer situations (Taylor et al., 2000). The funnel was placed 40 mm above the collecting beaker, and the beaker was placed on a measuring scale. The nozzle was fixed onto the funnel and the shutter mechanism was used to prevent any premature flow from the funnel. 100 g of the sample powder was transferred to the funnel and the test was started 30 seconds after the transfer (this facilitated floccule formation). The weight was recorded for the samples with respect to the time in triplicates (n=3) for each nozzle diameter (10, 15, and 25 mm) and samples (PA 12 (reference), granules, and drug). Time versus weight curves were constructed for the processed granules and the properties were compared with PA 12.

#### 2.4.5 Angle of repose

A 100 mm circular test platform together with a digital height gauge having a range of 0-300 mm and an accuracy of 0.03 mm was used (BEP2, Copley Scientific Limited, Nottingham, UK). The test platform had a protruding outer lip in order to retain a layer of the powder upon which the cone was formed. The surplus powder was collected in a tray below the test platform. The nozzle (10 mm nozzle for the angle of repose) of the funnel was placed 75 mm above the test platform, and the nozzle was secured using the shutter mechanism. 100 g of the sample (drug, and granules) were placed in the funnel, and the shutter was moved gently but rapidly to allow the powder to flow. The powder formed a conical on the test platform and started overflowing. The sample was allowed to overflow until the pile height was observed to be constant, this was protocol was repeated thrice (n=3). The height of the powder cone was measured using the digital height gauge and the diameter of the cone was 100 (diameter of the platform was 100 mm). Equation 2 was used to calculate the angle of repose.

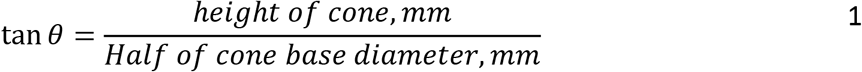

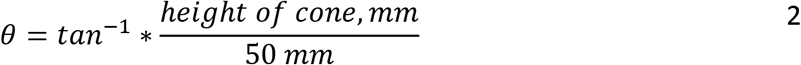

### 2.5 Laser sintering 3D printing

Granules (25% w/w) were mixed with Kollidon^®^ VA 64 (72% w/w), and candurin (3% w/w) (this mixture composition here on out will be referred to as PM-II). PM-II was used as the feedstock for the PBF based LS 3D printing process. This powder batch was introduced to the feed region of the benchtop LS 3D printer (Sintratec kit, Sintratec, Switzerland) equipped with a 2.3W 455nm laser. A powder batch of 150g was used for each build cycle. For the system set up the CAD file with twelve printlet having 5 mm height and 12 mm diameter were loaded on to the Sintratec central software, the coordinates of which have been depicted in fig. 2. Moving forward the layer height was set to 100μm, with the number of perimeters set to 1, and the perimeter offset set to 200μm. The Hatching offset was set to 120μm, and the hatch spacing was set to 25μm. After setting up the print parameters, the printing conditions were established where the chamber temperature was set at 80°C and the surface temperature was maintained at 105°C, which are both below the glass transition point of the polymer (>120°C) and the melting point of the drug (>160°C). Chamber temperatures maintained close to or higher than the surface temperature have been observed to form agglomerates and caused fusion of the blend in the feed region which ultimately leads to print failure [11]. The laser speed was maintained at 50mm/s. These process parameters were maintained for each build cycle and the build cycle was repeated thrice each time with a virgin powder batch to prevent possible degradation of the drug substance. Each manufacturing lot composed of twelve tablets and all the tablets were tested for their weight, and dimensions using a calibrated vernier caliper to evaluate the repeatability of the AM process. Moreover, the tablets from each printed batch were tested for hardness (n=3) (using a TA-XT2 analyzer (Texture Technologies Corp, New York, NY, USA)), disintegration time (n=3), dissolution studies (n=3), and other solid-state characterization techniques. Using the observed dimensions of the tablets, their volume was calculated using equation 3 where ‘r’ is the radius and ‘h’ is the height of the tablets. The density was then calculated using the volume and the weight of the tablets using equation 4. HME reference was manufactured using PM-II at 165°C instead of 155°C employing the same setup as described in fig. 1.

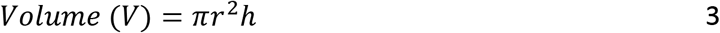

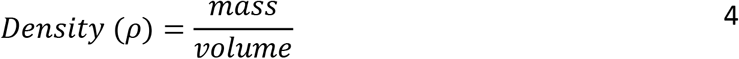

**Figure 2.**
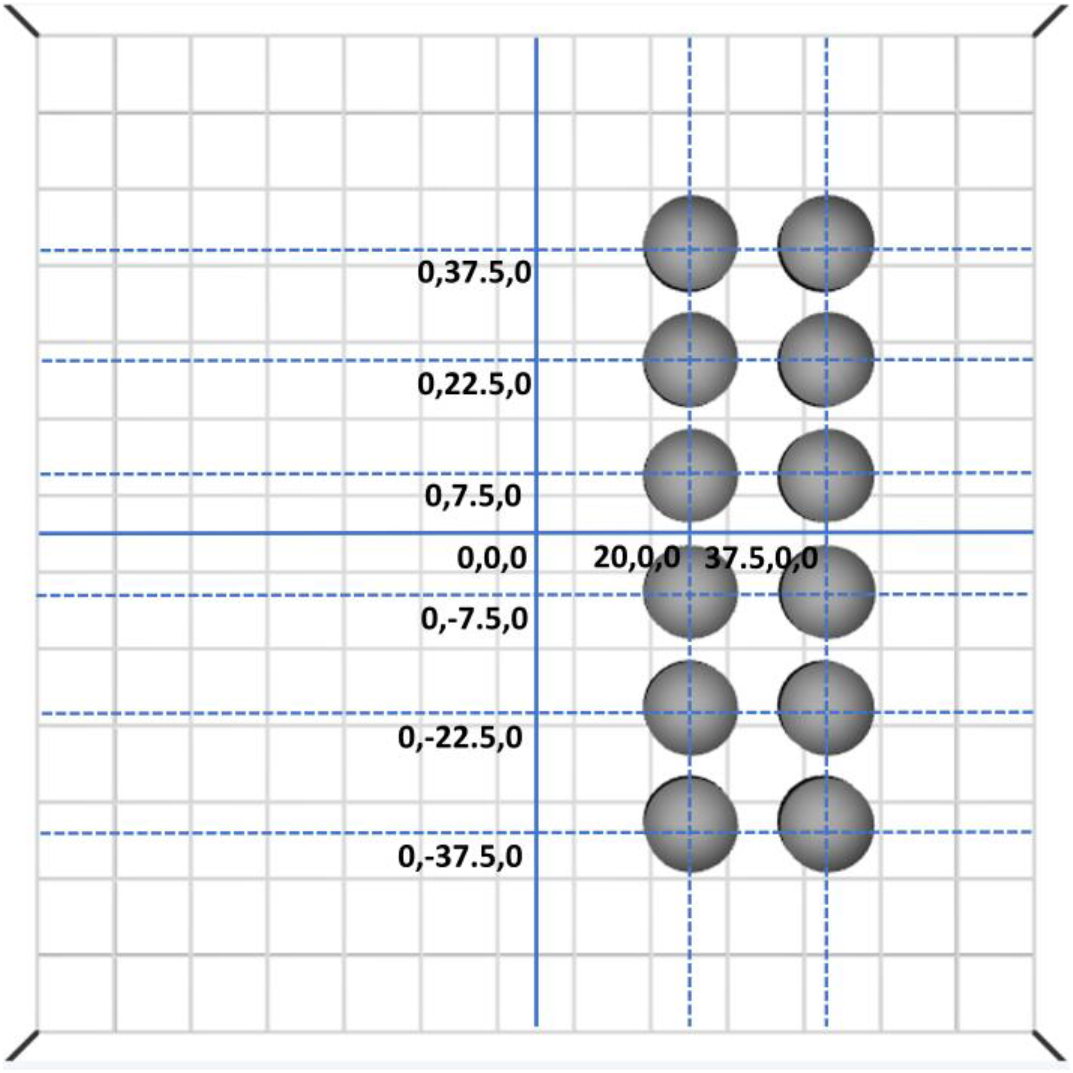
Positions of the parts (referred to as printlet or tablet in this manuscript) in the build platform with a maximum build volume 110 × 110 × 110 and a recommended build volume of 90 × 90 × 90 (units are in mm).

### 2.6 Solid-state characterization

#### 2.6.1 Powder X-ray diffraction (pXRD)

To investigate the solid-state of IND, PM-I, granules, Kollidon^®^ VA 64, PM-II, 3D printed tablets, and hot-melt extruded samples pXRD analysis was conducted. 100-150mg of the samples were dispensed onto the sample cell, the surface was flattened using a glass slide, the excess powder was discarded, and these cells were placed on the sample holders. The samples were then analyzed using a benchtop pXRD instrument (MiniFlex, Rigaku Corporation, Tokyo, Japan). The measuring conditions were set to a 2θ angle from 5 to 60 degrees, a scan speed of 2 degrees/minute, scan step of 0.02 degrees, where the resultant scan resolution observed was 0.0025. Moreover, the voltage and the current for the analysis were maintained at 45V and 15mV, respectively. The data were collected and plotted as a graph of intensity versus 2θ.

#### 2.6.2 Modulated differential scanning calorimetry (mDSC)

To investigate the presence of crystallinity or degradation mDSC analysis of IND, PM-I, granules, Kollidon^®^ VA 64, PM-II, 3D printed tablets, and hot-melt extruded samples were conducted (DSC Q20, TA^®^ instruments, Delaware, USA). The observations from the analysis were also used to determine the temperatures for the LS-AM process as well as HME processing. Samples weighing 5-15mg were dispensed in standard aluminum pans (DSC consumables incorporated, Minnesota, USA) using a microbalance (Sartorius 3.6P microbalance, Göttingen, Germany) and sealed using standard aluminum lids. The analysis was conducted from 60°C to 200°C, where the ramp rate was set to 5°C/minute, and modulation of ±1°C every 60 seconds. The collected data were analyzed by developing temperature (°C) versus reverse heat flow (mW) plots.

#### 2.6.3 Fourier transform infrared spectroscopy (FT-IR)

FT-IR provides insight into the post-processing interactions between different functional groups present on the components. FT-IR analysis was used to investigate the changes in the IND spectrum after each process and the interactions between IND and other components (iS50 FT-IR equipped with a SMART OMNI-Sampler, Nicolet, ThermoFisher Scientific, Waltham, Massachusetts, USA). A sample of 20-25 mg of IND, excipients, PM-I, granules, PM-II, LS printed tablets, and extruded filaments (powdered for the analysis) were dispensed on the sample holder and their % transmittance was measured using a range of 3100-700 cm^−1^. The resolution of the test was set to 4 cm^−1^ with 64 scans per run. To ensure the absence of contamination from previous samples, the cell was cleaned using isopropanol and the background spectrum was collected between each sample. The raw data were translated into spectra which were then investigated for intermolecular interaction, stability, and the solid-state of the samples using OMNIC™ series software (ThermoFisher Scientific, Waltham, Massachusetts, USA).

#### 2.6.4 Raman mapping

Raman surface mapping was conducted for the pure drug, the physical mixture, as well as the granules to evaluate the changes in the inelastic scattering between the samples and also check the drug distribution on the surface of the sample using an iS™ 50 Raman module (ThermoFisher Scientific, Waltham, Massachusetts, USA) equipped with an Indium Gallium Arsenide (InGaAs) detector, and an XT-KBr beam splitter. The samples were loaded on the sample holder and the sample surface was focused on using the associated microscope and camera, three different zones were analyzed for each sample to investigate the difference in the intensity of their spectrum which could be an indicator of the drug distribution throughout the sample. The power was set to 0.50W, the spectra were collected from 100-4000cm^−1^ with a resolution of 4cm^−1^, and the number of runs was set to 32 to reduce the noise. The data were collected as shifted spectrum and were plotted against the observed intensity.

### 2.7 HPLC method of analysis

The HPLC method was adopted from a previously conducted study [36]. IND was estimated using reverse phase-high performance liquid chromatographic (RP-HPLC) analysis (Agilent 1100 series, Agilent Technologies, Santa Clara, California, USA). A 250mmx4.6mm, 5μm particle size, stainless steel C-18 column (Nucleosil^®^100-5C18 (Suppleco series), Millipore Sigma, Burlington, Massachusetts) was used for the analysis. 0.2% o-phosphoric acid with acetonitrile (30:70) was used as the mobile phase. The flow rate and the injection volume were set to 1.2mL/min and 5μL, respectively. The retention time (R_T_) was observed to be 4.8 min and hence the run duration was maintained at 6 min. Indomethacin was detected using a UV-vis detector (Agilent 1100 series, Agilent Technologies, Santa Clara, California) at a wavelength of 237 nm. A calibration curve ranging from 0.5-8μg/mL was used for the quantification of indomethacin (R^2^=0.999).

### 2.8 *In vitro* release testing (IV-RT)

The performance of the manufactured granules and the 3D printed tablets were tested against pure crystalline IND, PM-I, PM-II, and HME samples using *in-vitro* release testing. For this pH-shift dissolution test, the aforementioned samples (n=3) were introduced into 750mL of HCl-KCl buffer (pH 2, 0.1M, hydrochloric acid-potassium chloride buffer) for 2 hours using a 900mL vessel. 150mL of phosphate buffer (pH 6.8, 0.1M) was then introduced to each vessel shifting the pH from 2 to 6.8, with a final volume of 900mL. It is important to note that the phosphate buffer should be prepared for a final volume of 900mL, hence the aforementioned 150mL volume is the concentrated buffer. The dissolution of the samples was tested at the final pH of 6.8 for an additional 2 hours. The test was conducted using a Paddle type assembly (USP type II). The test was conducted using a standard dissolution apparatus (Vankel VK 7000, Agilent Technologies, Santa Clara, California, USA) at 37.5°C, and the paddles were maintained at 50RPM throughout the study. Sample media of 1mL were drawn using 0.2μm polyethersulfone filters (VWR International, Radnor, Pennsylvania, USA) with an autosampler (Vankel VK 8000, Agilent Technologies, Santa Clara, California, USA) at predetermined time points. The sample volume was replaced with 1mL fresh buffer (HCl-KCl/Phosphate buffer) to maintain the volume in the vessels. Acetonitrile was used to dilute (2-fold) the drawn samples and the previously described method of analysis was used to quantify the API in the samples.

## 3. Results

### 3.1 Granulation and in-line monitoring

The hot-melt extrusion-based granulation process was developed using a modified screw design with mixing zones throughout the barrel to induce shear mediated melting, mixing, and absorption of the drug on to the inorganic excipients, and the process was monitored using UV-Visible reflectance probe placed in the ‘zone 6’ of the barrel where the drug was expected to have completely converted into its amorphous form under the right conditions. The granulation process was monitored using the process analytical tool (PAT) to monitor the stability of the process and to monitor the amorphous conversion of IND. CIELAB yellow-blue color space coordinate (b*), and yellowness index (E313-00 YI) were monitored to exploit indomethacin’s unique property as in its crystalline γ-form it exhibits a pinkish white appearance, whereas on amorphous conversion its color changes to yellow [34]. Fluctuations in these aforementioned parameters were expected to indicate a combination of crystalline and amorphous indomethacin in zone 6. These fluctuations can be observed in fig. 3 (A, B, and C) which depict incomplete amorphous conversion of indomethacin which was also observed in the pXRD patterns for the collected samples at these temperatures i.e., 140-150°C. The other monitored parameters such as the custom selected wavelength (600-700 nm) which depicts the average response between 600 and 700 nm, wavelength of maximum reflectance over the measured region (PWL), and reflectance value at the PWL (Peak) were utilized to monitor the stability of process as the fluctuations in these parameters were expected to be a result of improper mixing which would eventually lead to a content uniformity with a broader standard deviation. All the monitored parameters stabilized at 155°C as seen in fig. 3 D. The samples collected at 155°C after the process was stabilized were found to be amorphous on pXRD analysis hence the batches for LS 3D printing were manufactured at 155°C and the process was monitored using the aforementioned PAT and parameters.

**Figure 3.**
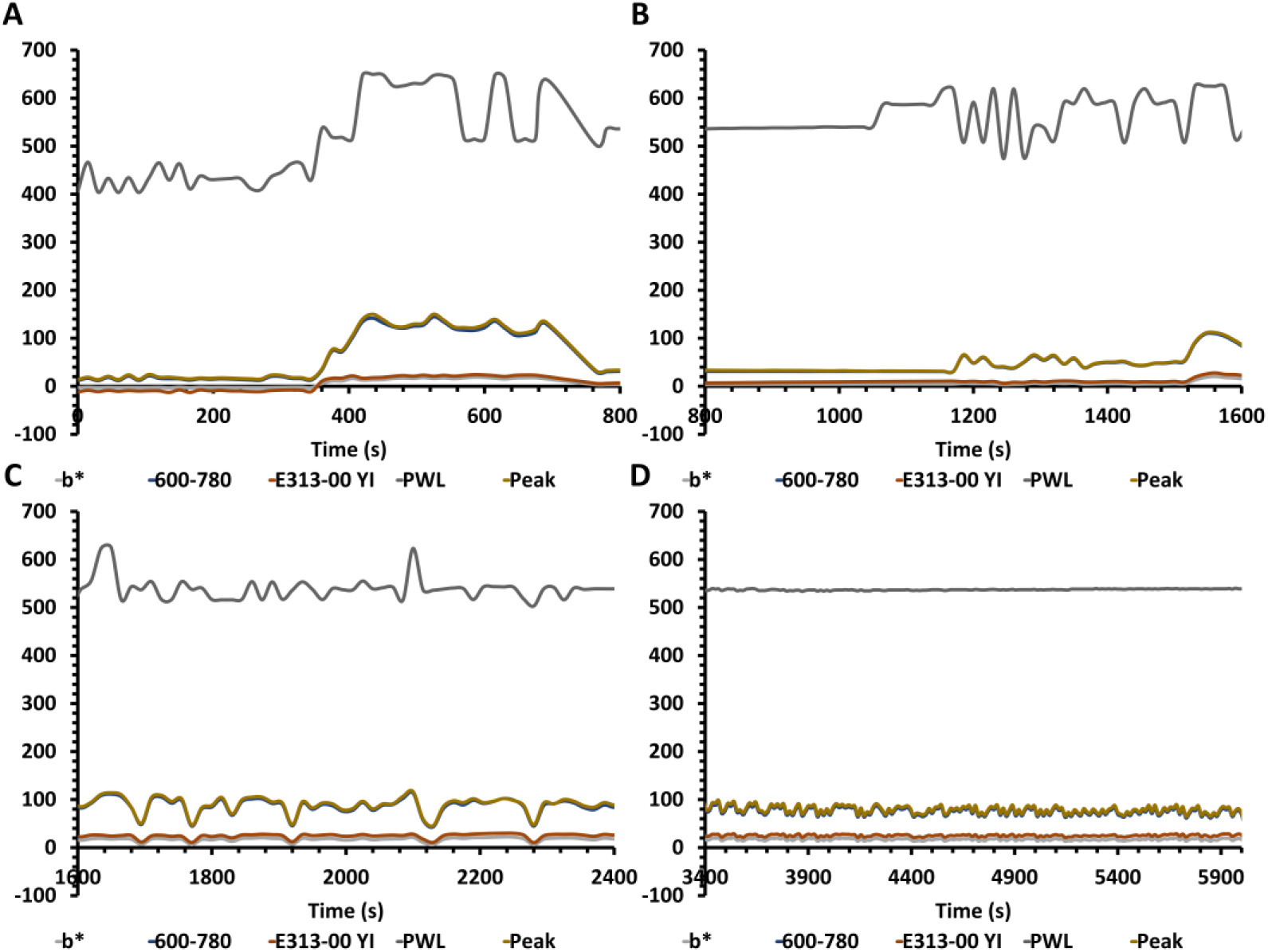
Process monitoring of CIELAB yellow-blue color space coordinate (b*), custom selected wavelength (600-700 nm), yellowness index (‘E313-00 YI’ which is supposed to trend with b*), the wavelength of maximum reflectance over the measured region (PWL), and reflectance value at the PWL (Peak) with UV-Visible reflectance probe at different extrusion temperatures at A) at 140°C B) at 145°C C) at 150°C D) at 155°C.

Moreover, the drug content uniformity conducted by sampling the PM-I from three different regions for the three manufactured batches was found to be 101.17±5.64%, 103.68±7.64%, and 100.25±6.67% respectively, whereas the granules collected after HME processing were found to have content uniformity of 100.39±0.51%, 99.86±0.93%, 99.87±0.85% respectively.

### 3.2 Bulk properties testing

Previous research has shown that indomethacin has extremely poor flow properties by the means of Angle of Repose (AOR) and Hausner’s ratio [32]. On inspecting indomethacin drug crystals using digital microscopy (fig. 4 A-1), polarized light microscopy (fig. 4 B-1, fig. 4 C-1), and scanning electron microscopy (fig. 4 D-1) it was observed that indomethacin crystal have uneven and rough surfaces which highly contribute to the observed poor flow properties.

**Figure 4.**
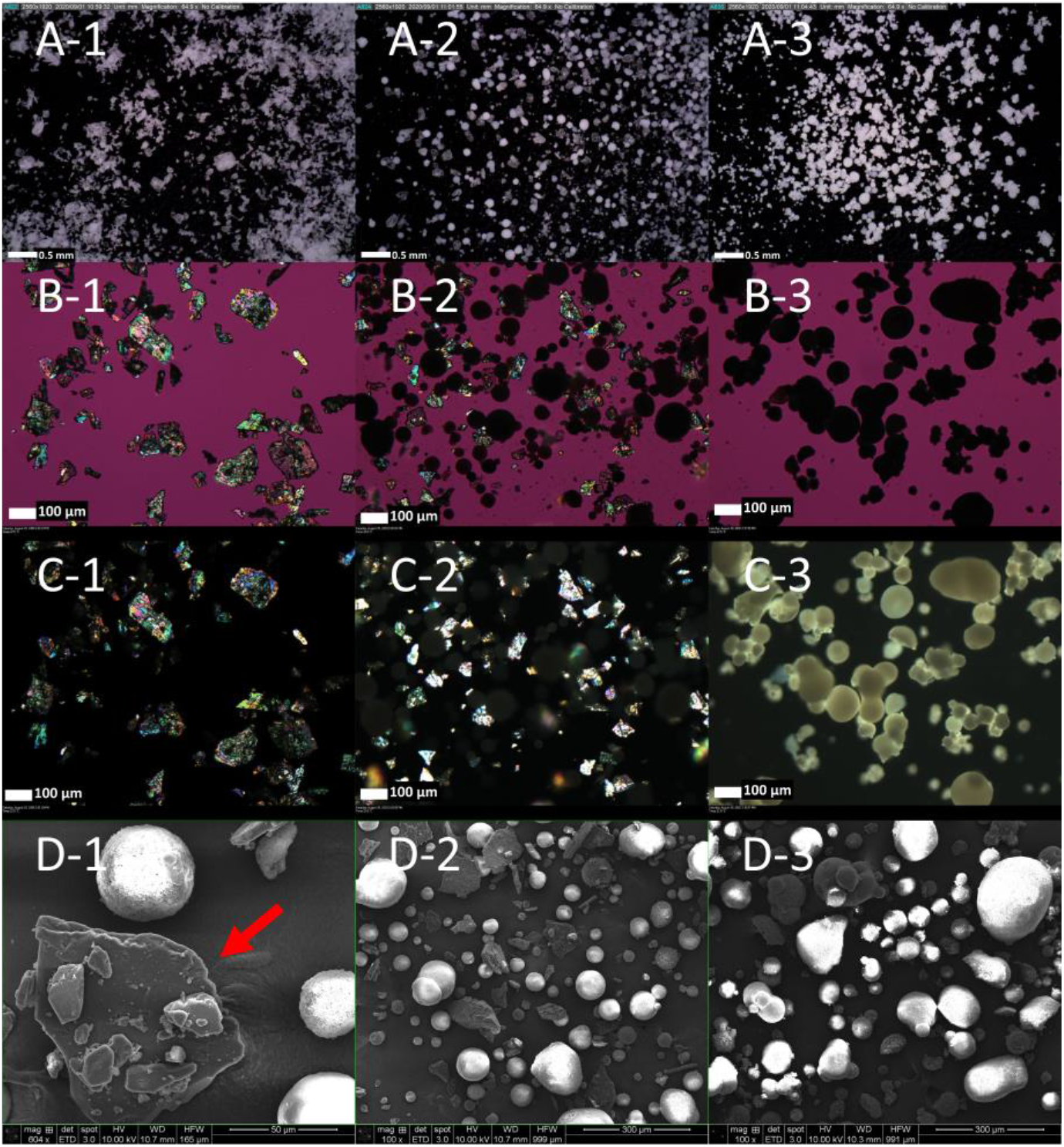
A) Digital microscopy images, A-1) Indomethacin crystals, A-2) Physical Mixture-I, A-3) Processed granules; B) Polarized light microscopy (530 nm compensator) images, B-1) Indomethacin crystals, B-2) Physical Mixture-I, B-3)Processed granules; C) Polarized Light Microscopy (dark mode), C-1) Indomethacin crystals, C-2) Physical Mixture-I, C-3) Processed granules; D) Scanning Electron Microscopy, D-1) highlighted Indomethacin crystal, D-2) Physical Mixture-I, D-3) Processed granules.

The AOR analysis of IND was conducted as a reference to the manufactured granules and it was found to have a θ value of 58.2±0.06 degrees, which is classified as ‘very poor’ as per the AOR reference table. Furthermore, the flow through orifice analysis of pure IND could not be conducted as it clogged the orifice and was observed to have no flow throughout the test.

Moving forward to the physical mixtures depicted in fig. 4 (A-2, and D-2) which show the discrepancies between the sizes of the drug crystals and the inorganic excipients further shed light on the broad standard deviations observed for PM-I. Physical mixtures having components with different densities tend to segregate when exposed to prolonged vibrations during mass transfer processes. PM-I depicted irregular trends because of repetitive clogging of the orifice and non-reproducible results when exposed to flow through orifice analysis and hence these trends were not depicted in the results section. Further, fig. 4 (B-2 and C-2) depict the PLM images of PM-I under the microscope which highlight the presence of crystallinity along with other amorphous inorganic excipients in the blend which do not show any birefringence and hence are seen as dark spots in the images.

The granulation technique was designed to break down the crystalline drug and further facilitate the absorption of the drug onto the surface of the inorganic excipient. During post-processing, as expected, it was observed that the drug crystals completely disappeared (fig. 4 (A-3, B-3, C-3, and D-3)). Moreover, the birefringence observed in fig. 4 (B-2 and C-2) was not present post-processing in fig. 4 (B-3 and C-3) indicating complete amorphous conversion of the drug from its crystalline counterpart [37]. It can also be seen in fig. 4 (A-3, and D-3) that the granules have a smooth and round surface which can translate to better flow properties as compared to PM-I. The AOR of the extruded granules were observed to have a θ value of 29.49±1.24 degrees, which is classified as ‘excellent’. On conducting the flow through orifice test for the granules and PA-12 (LS reference material), it was observed that the granules depicted a reproducible trend just like PA-12 with an R^2^ value of ‘>0.9’ for all three orifices (10, 15, 25 mm). Although both the materials demonstrated different weight/minute values, the rationale of this test was to stimulate the flow in a hopper or other mass transfer situations and inspect if the flow follows a reproducible trend. The granules observed excellent trends and no significant deviation as seen in fig. 5.

**Figure 5.**
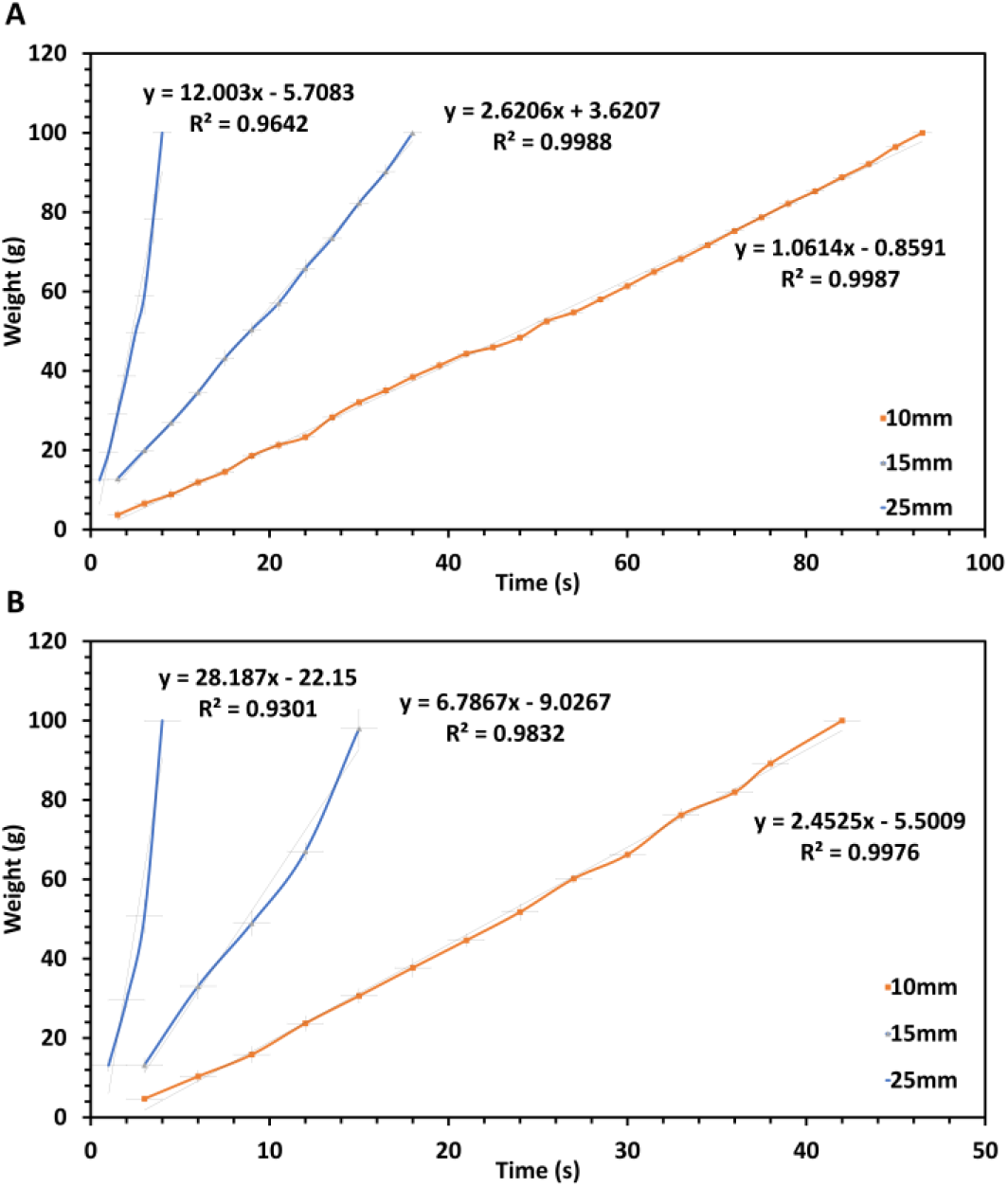
Flow-through orifice ‘weight versus time’ plots for three different orifice diameters (10, 15, 25 mm) A) Extruded granules B) PA-12 (LS reference material).

**Figure 6.**
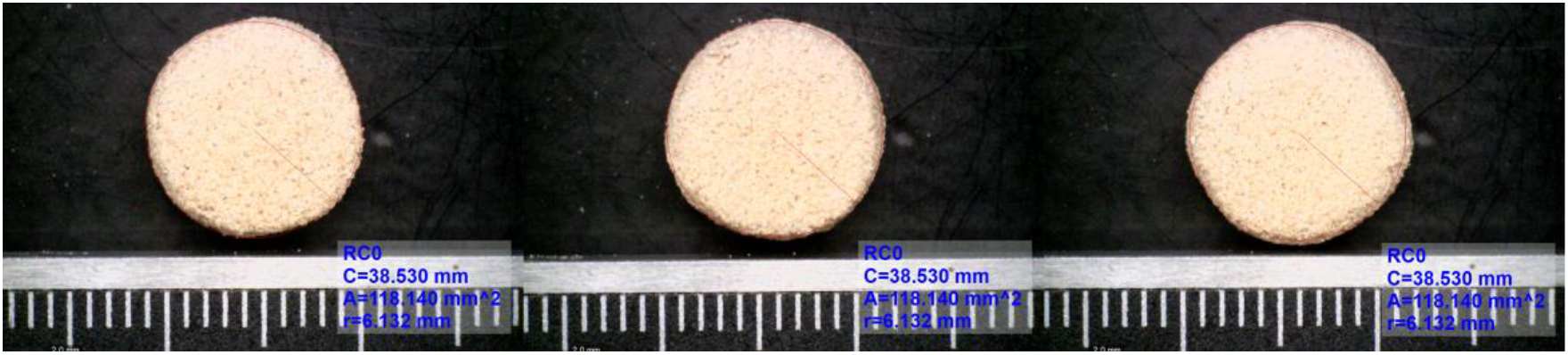
Morphology of LS 3D printed indomethacin tablets using HME based granulation technique.

### 3.3 3D printing process

During the LS 3D printing process, 100μm thick layers of the PM-II from the feed region were spread onto the build platform where the particles were sintered together utilizing a laser as per the design of the tablets. During the build cycle, the build surface maintained its smooth texture as the PM-II exhibited excellent flow properties as expected and there were no print failures or defects experienced during the printing process after the processing parameters were fixed. The twelve tablets manufactured per build cycle were isolated from the build chamber and de-dusted to remove the part cake using a nozzle-based air dispenser. The tablet dimensions were measured and are depicted in Table 1. It was observed that all the tablets were printed as per the dimensions of the designed tablet with an insignificant standard deviation and batch to batch variation, moreover, the deviation in the weights was within the defined range as per the United States Pharmacopoeia (USP).

**Table 1.**
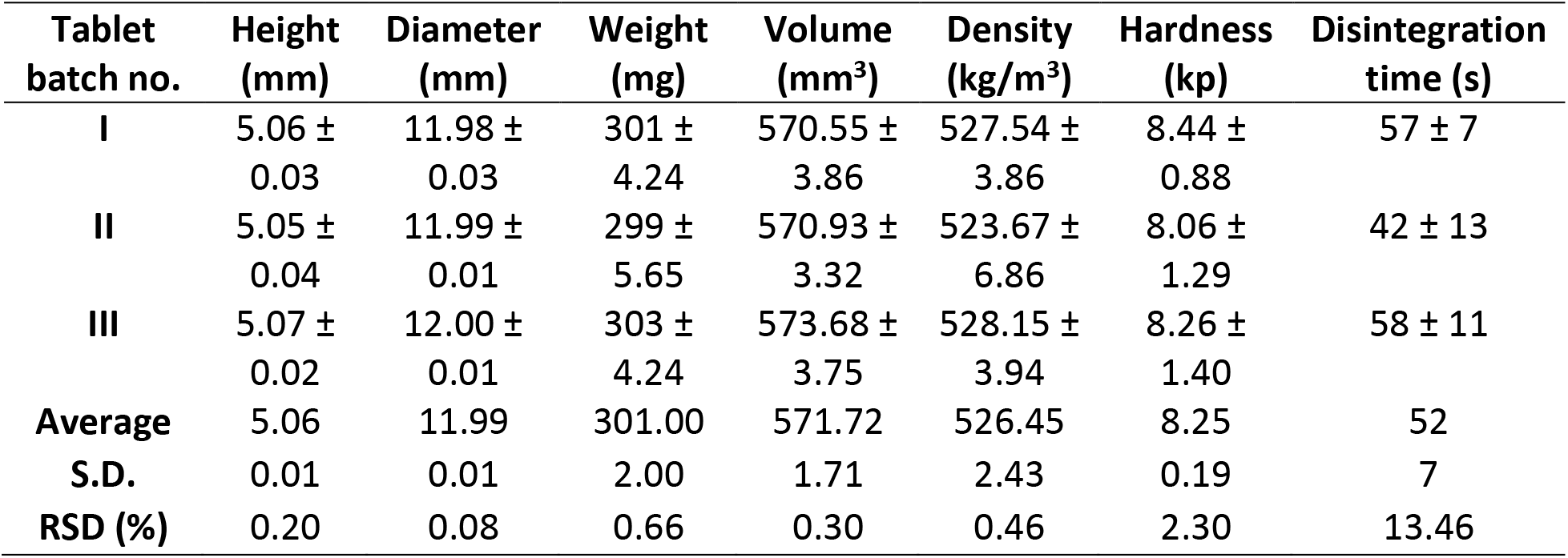
Quality control characteristics of the manufactured 3D printed tablets.

The target weight for these tablets was 300mg and as per USP 7.5% weight variation is acceptable for tablets with the mentioned weight, therefore a deviation of ±22.5mg would be considered acceptable, although the observed deviation was extremely narrow.

The volume and the density were calculated from the dimensions and the weight of the tablets, the density of a single layer can be used to predict the dimensions for a tablet with the desirable weight. The hardness of the tablets was also found to be reproducible. The inherent property of LS 3D printing allows the manufacturing of porous tablets. Even though the tablets have sufficient hardness, they disintegrate in less than 60s when they come in contact with aqueous buffers.

### 3.4 Solid-state characterization

X-ray diffraction has found applicability as one of the most reliable and straightforward techniques for qualitative and quantitative analysis of crystallinity. One of the major aims of this research was the complete amorphous conversion of the drug post HME processing and LS 3D printing. XRD was used as a tool to optimize the manufacturing temperature and to inspect the solid-state of the optimized granules and manufactured tablets. For the pure drug, all the characteristic peaks of IND (γ-form) were observed (fig. 7) at 11.5, 19.4, and 21.7 two-theta (2θ) degrees, which comply with previous reports [38].

**Figure 7.**
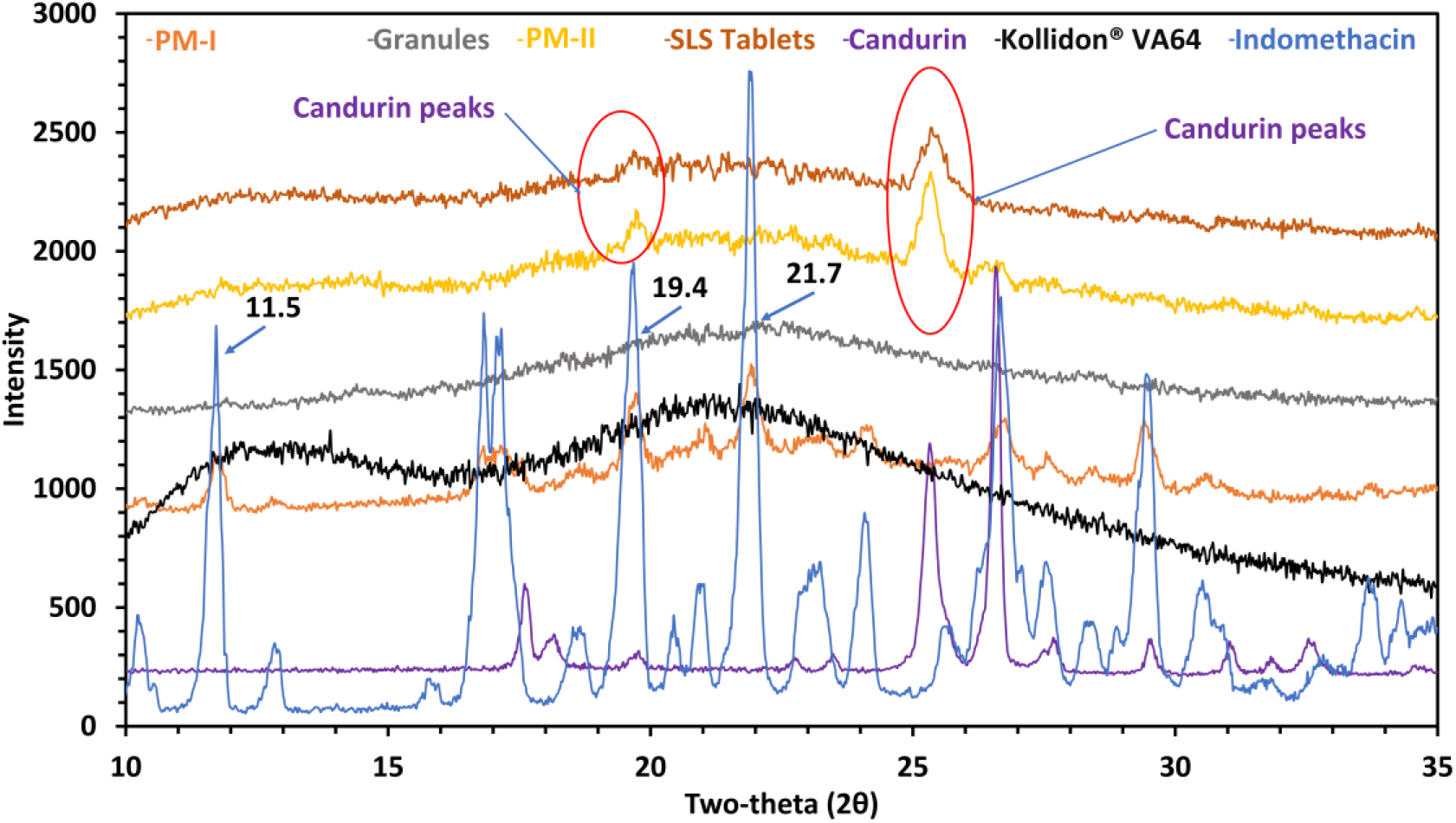
Powder X-ray diffraction analysis of indomethacin, excipients, physical mixtures, extruded granules, and 3D printed tablets.

Polymers such as Kollidon^®^ VA64 are usually semi-crystalline or amorphous and hence do not depict any characteristic peak on pXRD analysis. Although the trace crystallinity of the polymers can be observed using techniques such as wide-angle X-ray scattering (WAXS), this technique was not used for this research as we wanted to focus on the solid-state of the drug and not the polymer. All the characteristic peaks of IND were observed in the PM-I before it was processed using twin-screw extrusion. Post-processing all the peaks disappeared which indicates a change in the solid-state of the drug from its crystalline to its amorphous form. Further, there are two small peaks at 19.8 and 25.2 two-theta (2θ) degrees, out of which one overlaps with the characteristic peak of IND at 19.4 two-theta (2θ) degrees. These peaks belong to Candurin^®^ which is the photo-absorbing species required for the successful sintering of PM-II and hence should not be confused with the presence of trace crystallinity in the LS 3D printed tablets [39]. This can further be proven by the mDSC results depicted in fig. 8. The thermal analysis of all the samples was conducted to verify the absence of crystallinity in the granules and the LS 3D printed tablets, as the drug in its amorphous form does not exhibit a melting endotherm. The melting endotherm of IND was observed in both the pure drug and the PM-I sample which represents the melting point (T_m_ = 160.4°C) of IND (for the γ-form). Moreover, mDSC also revealed the presence of polymorphism in the IND drug crystals. Indomethacin has a glass transition (T_g_) of 42°C (not shown here), after which it recrystallizes to its metastable α-form (fig. 8) close to 100°C which shows a melting endotherm at 152-154°C as seen in fig. 8, whereas the more stable γ form melts at 160°C on the application of heat energy, this transition has been depicted by the means of dotted lines in fig. 8 [40].

**Figure 8.**
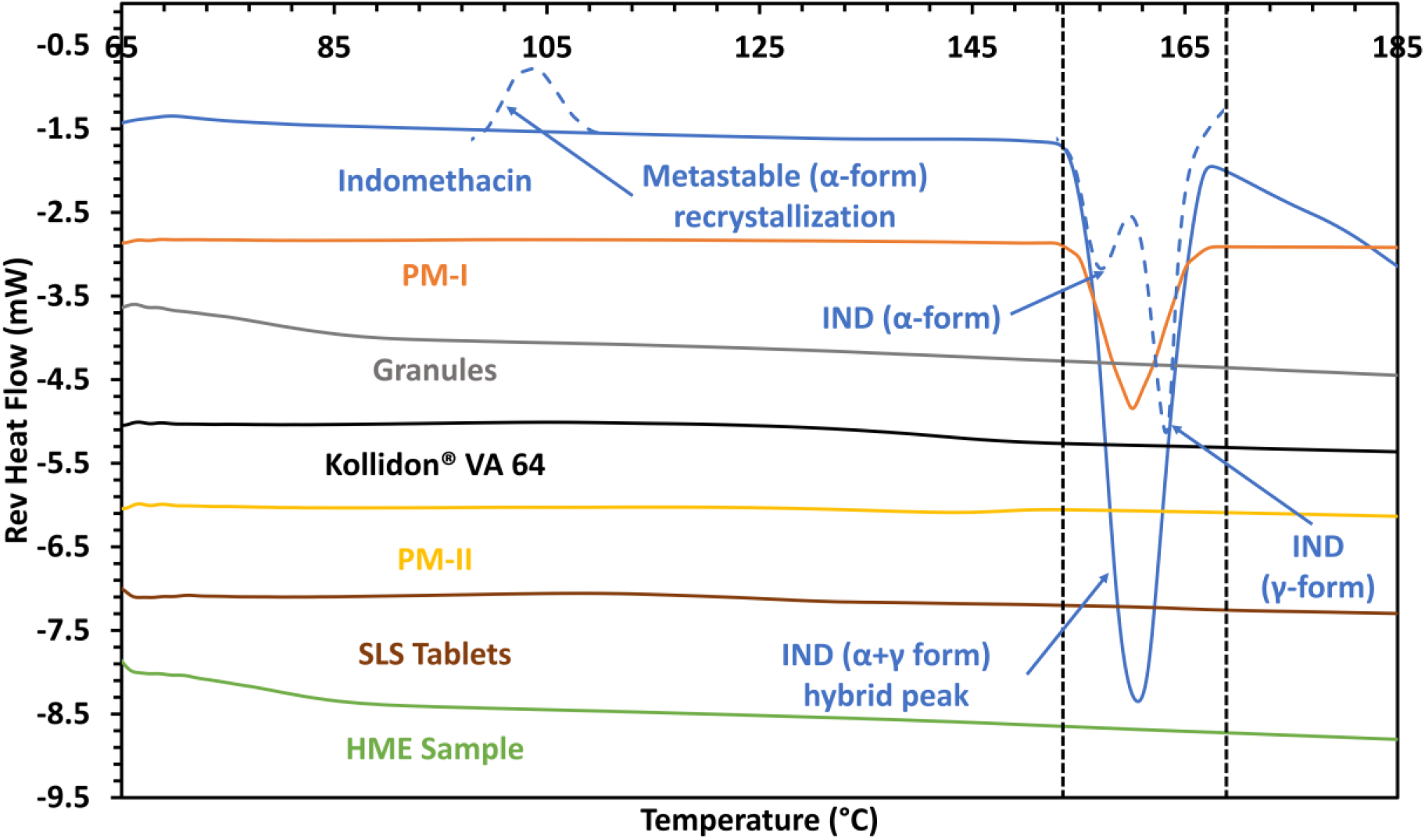
Modulated differential scanning calorimetry of IND, excipients, physical mixtures, extruded granules, and 3D printed tablets.

Observations made from the pXRD data were further bolstered by the DSC data where no melting endotherms were observed for the granules or the 3D printed tablets. These results suggest that the solid-state of the drug was amorphous in the granules and tablets. Furthermore, it was also observed that IND starts to degrade on increasing the temperature over its T_m_. It has been previously observed that IND starts to degrade over its T_m_ when exposed to the conditions for an extended amount of time and gets converted to decarboxylated IND [41].

Hence, the HME conditions selected (155°C) were safe for processing IND as it was below the T_m_ and the degradation temperature (T_d_) of the drug. Further FT-IR studies conducted on all the samples provided further insight into the solid-state as well as the interactions between IND and other ingredients in the formulation. FT-IR samples of IND had confirmatory peaks at 2926.6 cm^−1^ (O–H stretching vibration), 1717.0 cm^−1^ (C=O stretching of carboxylic acid dimer), 1691.5 cm^−1^ (C=O stretching of benzoyl group), 1307.7 cm^−1^ (C–O), and 1068.0 cm^−1^ (C–Cl) which are represented in fig. 9.

**Figure 9.**
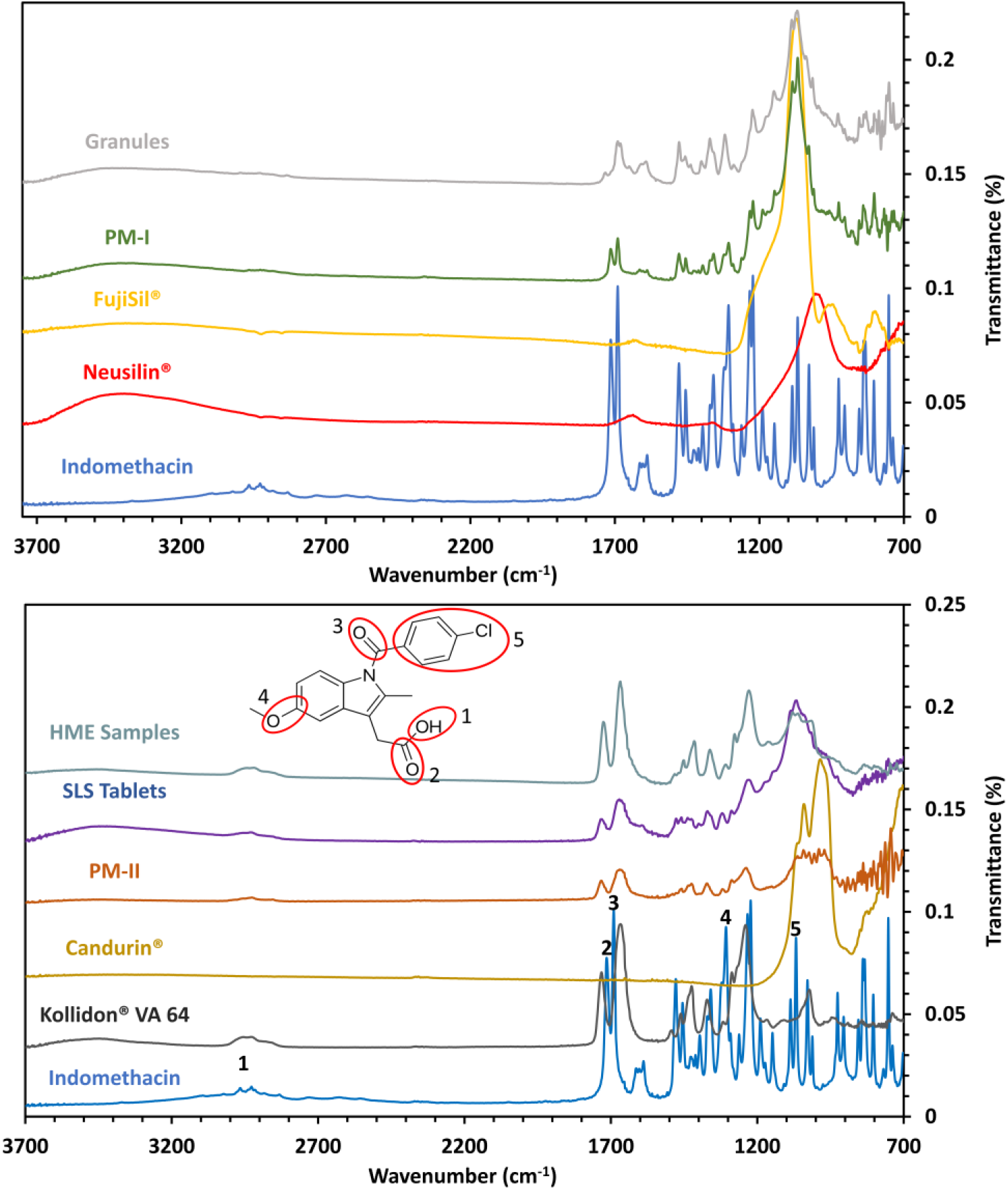
Fourier transform infrared spectroscopy of IND, excipients, physical mixtures, extruded granules, and 3D printed tablets.

The carbonyl groups part of amides (associated with nitrogen atoms) usually exhibits peaks at lower wavenumbers (1640 cm^−1^) [42]. Although, the group being an indole ketone, experiences a reduction in the contribution of the mesomeric effect (where nitrogen can donate its lone pair of electrons) in molecules as the nitrogen atom is part of the ring. The mesomerism is responsible for the lower wavenumbers observed for the typical amides and since the effect is absent in IND, the wavenumber for the benzoyl group in the γ-form is relatively higher [43]. Furthermore, the peaks at 1717.0 cm^−1^ are representative of the carboxylic acid cyclic dimer (hydrogen bonding between two sterically unhindered groups of carboxylic acid) [44]. The formation and existence of these dimers are crucial for the stability and re-crystallization of IND post amorphous conversion. The retention of Peaks 2 and 3 in all samples suggest that there was no chemical degradation or unwanted chemical reaction between IND and other excipients. The changes in the peaks post-processing were expected due to the amorphous conversion of the drug. Post amorphous conversion the benzoyl carbonyl peak shifted from 1691 cm^−1^ to 1684 cm^−1^ as in the crystalline γ form. The carbonyl group is stabilized by the hydrogen bonds from the hydroxyl group of the neighboring IND molecule. The peak for the carboxylic acid dimer shifted from 1717.0 cm^−1^ to 1733.0 cm^−1^.

As per previous reports post amorphous conversion, IND exists in two forms i.e. cyclic dimers (1706 cm^−1^) and non-hydrogen bonded carboxylic acid (1735 cm^−1^) [43]. In this study, post-processing IND was observed to dominantly exist in just one state i.e., non-hydrogen bonded carboxylic acid, which might be because of the surfactant and the inorganic excipients as well as their contributions in stabilizing IND in its amorphous form. The absence of dimers might significantly reduce the rate of re-crystallization for the granules although the solid-state stability studies were beyond the scope of this current research and are being conducted as an extension to it. The FT-IR of the 3D printed tablets and the HME samples did not depict strong drug peaks for two reasons, firstly the presence of Kollidon^®^ VA64 overshadowed the drug peaks, and secondly, the drug and the polymer post-processing were expected to form an amorphous solid dispersion. Although there was a resemblance between the FT-IR of the HME and the LS 3D printed samples which can be attributed to their similar composition and intermolecular interactions.

Raman spectroscopy like FT-IR, demonstrated the peak shift from 1700 cm^−1^ to 1680 cm^−1^ which represents the benzoyl carbonyl stretching and hence does not give any information of the molecular associations however, it does provide some information on the influence by steric hindrance and tension of molecules imposed by hydrogen-bonded associations in dimmers in the γ form [42,45]. The sharpness and intensity of this band are associated with the long-range of the dimer chains and this is apparent in the Raman spectra of PM-I (fig. 10 B), in contrast, the peak reduces in intensity and sharpness in the processed granules (fig. 10 A) where the drug is in its amorphous form. Furthermore, the Raman spectra of IND in its amorphous state are expected to exhibit a broad band around 1679 cm^−1^ (1698 cm^−1^ in its crystalline form, fig. 10 D) which corresponds to the dimer associations discussed previously. Although the carbonyl peak from the carboxylic acid disappeared from 1698 cm^−1^ as seen in fig. 10 C, it appears as a weak broad peak at 1680 cm^−1^ which confirms that the dimers do not exist in excessive quantities in the granules. This peak shift is prominent, and the intensity is quantifiable when IND is converted to its amorphous form by other techniques such as quench cooling [44].

**Figure 10.**
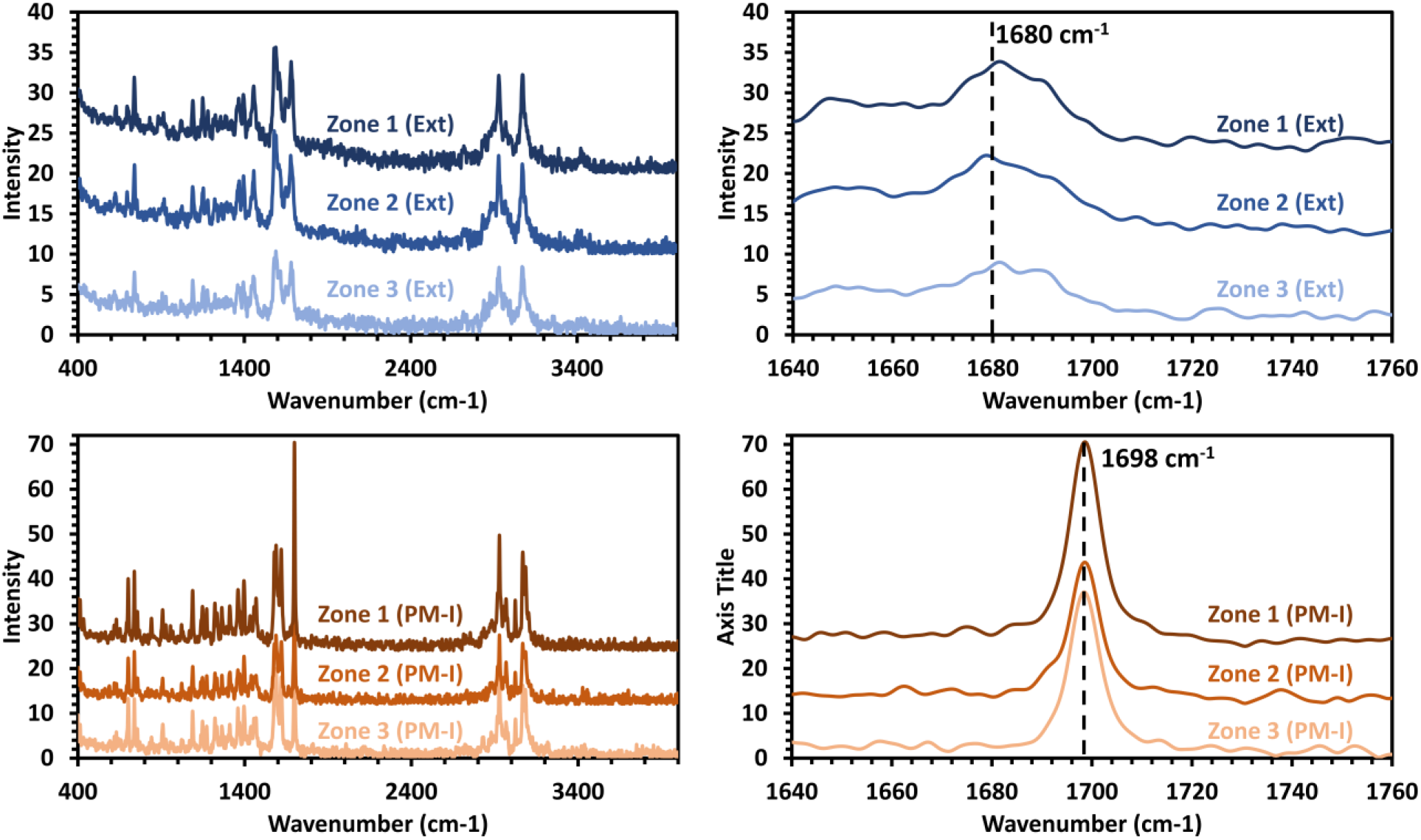
FT-Raman spectra of A) extruded granules B) Pre-extrusion physical mixture (PM-I) C) Shift in ‘carbonyl stretching’ region because of amorphous conversion post extrusion D) Carbonyl stretching region of crystalline IND in PM-I.

### 3.5 Performance testing

The *in vitro* release testing was performed using a pH shift protocol to simulate physiological conditions and contrary to the USP dissolution medium i.e., pH 7.2 phosphate buffer. The performance test was conducted in 250mL HCl-KCl buffer (pH 2.0) and then in 900mL phosphate buffer (pH 6.8). The rationale behind the pH shift dissolution study apart from mimicking the physiological condition was the pH-dependent solubility of IND. The drug is weakly acidic (because of the carboxylic group, pKa 4.5) and poorly soluble (because of the hydrophobic indole group) in nature [46]. IND is practically insoluble in acidic pH and has a solubility of 7-9μg/mL at 35°C in water, although it has a higher solubility in alkaline pH because of ionization of the molecule ^33^. To understand the impact of the granulation process, and further the LS 3D printing on the dissolution of the drug, it was important to conduct a pH shift dissolution test. Moreover, because of its pKa and hydrophobicity, and the amorphous nature of the drug in the samples of interest, it was important to expose it to the acidic conditions before exposing it to the relatively basic conditions. Weakly acidic and hydrophobic drugs like IND tend to undergo a solid-state transfer from their more reactive and less stable amorphous forms (higher chemical potential) to less reactive and more stable crystalline form (lower chemical potential) [47–49]. If the drug exists in its amorphous form in the formulation, it is crucial to study its stability in the acidic pH as in the physiological environment the formulation will be exposed to it first. Even if the formulation exhibits excellent dissolution at a relatively basic pH *in vitro,* but is unstable in acidic pH, there is a chance that it might not demonstrate a similar trend and be as effective as predicted due to recrystallization *in vivo*.

The amorphous form of the drug is of importance here as thermodynamically it has a higher chemical potential as well as reactivity and hence exhibits better and faster dissolution as compared to its stable crystalline counterpart which is less reactive and has a lower chemical potential due to the stability induced by its neighboring molecules [50]. This phenomenon is even stronger in this case where the IND molecules are stabilized by strong dimers which makes it difficult for the water molecules to break these bonds.

In the case of amorphous solid dispersions, hydrophilic polymers are used to break these intermolecular bonds between drug molecules which are originally in their crystalline form and forms new intermolecular interactions with the polymers or other excipients such as stabilizers and surfactants to prevent recrystallization [51,52]. For this formulation, the drug was absorbed onto inorganic carriers and was stabilized using polysorbate 80, moreover, the drug was found to be in its amorphous form. When exposed to an acidic environment (fig. 11 B), the pure drug crystals did not show any improvement in their solubility which was expected because of the above-discussed reasons. When the PM-I was exposed to pH 2, it demonstrated a slight improvement over the pure drug which is not significantly different from the pure drug, this slight improvement can be attributed to the presence of polysorbate 80 in the formulation which acts as a surfactant and facilitates the interactions between the drug and the medium by reducing the surface tension. Since the drug in PM-I was still in its crystalline form, the rate of dissolution was extremely slow. The performance of the granules on the other hand is significantly faster than that of the pure drug and the PM-I, although there seems to be a drop in IND solubility around 90-minutes into the dissolution studies which might be due to the recrystallization or precipitation of the dissolved drug. For this study, 0.2μ filters were used to make sure only the solubilized IND is estimated, and any recrystallized nuclei are filtered out. Moreover, the dissolution was faster than recrystallization and hence the drug concentration increased again on the 120-minute time point. The dissolution rate PM-II was observed to be similar to that of the granules. Furthermore, the LS printed tablets and the HME samples demonstrated the highest increase in the rate of solubilization as compared to the pure crystalline drug and the PM-I. The dissolution pattern of the HME samples and the tablets were also identical with little variability as both demonstrated around 4% drug release over a 2-hour dissolution test.

**Figure 11.**
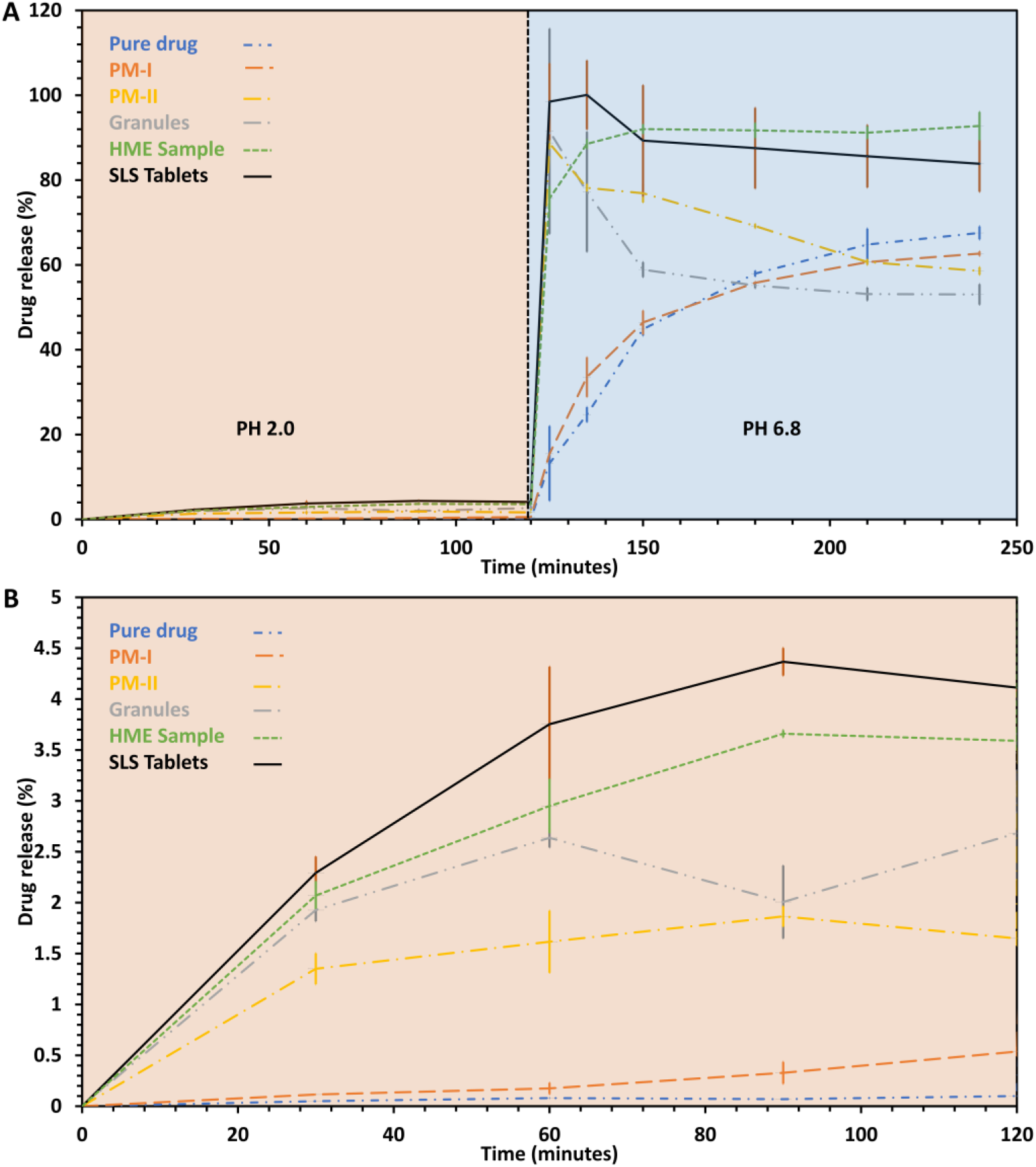
A) pH shift dissolution study for pure crystalline IND, PM-I, PM-II, Granules, LS 3D printed tablets, hot melt-extruded reference amorphous solid dispersion. B) Dissolution study for pure crystalline IND, PM-I, PM-II, LS 3D printed tablets, hot melt-extruded reference amorphous solid dispersion at pH 2.

When the pH of the dissolution medium was shifted from 2 to 6.8, the granules, PM-II, LS 3D printed tablets and the HME samples demonstrated >80% drug release in <5minutes. This rapid dissolution and completeness of the release of these formulations signified minimum to no recrystallization in the acidic pH. It can be seen that even though the dissolution rate of the pure drug and PM-I is faster as compared to their dissolution in the acidic pH, it is significantly slower than the other samples. The pure drug and the PM-I released only 60% of the drug over 2 hours. The drug release of the granules peaked at 91% and then depicted a rapid reduction in the drug concentration. This rapid reduction can be attributed to the recrystallization of the solubilized IND in the medium which was expected because even though the drug existed in its amorphous state in the granules which was responsible for its rapid dissolution, there was nothing in the formulation to maintain the drug in its solubilized state (less stable) and hence it recrystallized (more stable) in the medium. PM-II on the other hand peaked at 88% drug release and then started to recrystallize in the solution, although the recrystallization rate was slightly slower as compared to the granules which can be attributed to the presence of Kollidon^®^ VA 64 in the formulation. The dissolution profile of the LS printed tablets and the HME samples were still similar, where the HME samples peaked at 92% drug release (30 minutes after the pH shift) whereas the LS 3D printed tablets peaked at 100% drug release (5 minutes after the pH shift). Furthermore, the HME samples maintained the saturation of the drug in the medium throughout the dissolution study and the incomplete release can be attributed to the 2-hour exposure of the sample to the acidic pH and recrystallization of the free amorphous drug. Whereas, the LS 3D printed tablet demonstrated a 100% release within 5 minutes and a steep decline thereafter to 90% which can be attributed to the recrystallization of solubilized unstable IND in the medium. After the steep decline, the drug concentration for the tablets was maintained above 80% throughout the dissolution study which suggests ASD formation and stabilization by Kollidon^®^ VA64 during the LS 3D printing process.

## 4. Discussion

Indomethacin along with several other poorly water-soluble drugs have been used as models to demonstrated the versatility and applicability of material extrusion AM techniques in solubility enhancement and dosage form manufacturing [53,54]. Polymers such as polycaprolactone, polyethylene oxide, Affinisol™ HPMC HME 15LV, 100LV, hydroxypropyl methylcellulose K15M, hydroxypropyl methylcellulose acetate succinate, and poly(ethylene-co-vinyl acetate) (vinyl acetate 12 wt%) have been processed into filaments suitable for material extrusion-3D printing processes. Although, most of these FDM based processes were conducted over the melting temperature of the drug based on the polymer used. These high processing temperatures can lead to the generation of decarboxylated indomethacin as the drug starts to degrade over its melting point (i.e. 160°C) [41].

Considering the high processing temperatures employed in both hot-melt extrusion-based and FDM based processes, powder bed based AM techniques such as binder jetting and powder bed fusion can serve as alternatives for such thermolabile drugs. Although these processes have their challenges such as excipient selection and process repeatability. Furthermore, these powder bed-based techniques are useful in making porous structures and hence only found applicability for processing drugs such as levetiracetam [29] and acetaminophen [15], until we demonstrated the applicability of laser sintering 3D printing in the manufacturing of amorphous solid dispersion dosage forms for improved solubility of poorly soluble and thermolabile drugs such as ritonavir [11]. In our previous study, we encountered multiple printing-related issues leading to print failures which were mostly because of the powder bulk properties, which eventually led to weight variability and content uniformity issues in some of the formulations. Furthermore, processing aids were needed to ensure the proper flow of the physical mixture from the feed region to the build platform and consecutive build surfaces [11,17]. It was apparent that granulation principles used in conventional manufacturing to facilitate smooth mass transfer and content uniformity can also find applicability in these novel techniques although conventional granulation techniques might not be suitable because of the powder particle size (<100μ) requirement in these AM techniques [17,21]. In this research, we demonstrated an HME based granulation technique using a poorly water-soluble, thermo-labile drug, and inorganic mesoporous carriers, where the process was conducted below the melting point of the drug under investigation. These granules demonstrated excellent flow properties, stabilized the drug in its amorphous form, and also demonstrated immediate dissolution in comparison to the crystalline drug. Indomethacin solid dispersions with inorganic excipients such as mesoporous silica have been manufactured previously using solvent evaporation techniques such as rotary evaporation, spray drying, and vacuum drying [55–57] where hydroxypropyl methylcellulose (HPMC) and Kollicoat^®^ IR were used as precipitation inhibitors (PIs) combined with mesoporous silica nanoparticles (MSNs) as carriers. The present study manufactured granules with multiple mesoporous inorganic excipients, using a solvent-free, continuous process with in-line monitoring. Furthermore, these granules were used for LS based 3D printing process which further improved the performance of the formulation. It was observed that there was an increase in recrystallization stabilization with the presence of the polymer. This enhanced stability has been attributed to the increase in viscosity because of the molecular weight of the polymer, which in turn slows down the mobility of these systems [58]. However, in the dissolution medium IND interacts with Kollidon^®^ VA 64 by the formation of hydrogen bonds between the carboxylate group in the drug and the carbonyl group on 2-pyrrolidone in the polymer [59] which can explain the maintained saturation in the hot-melt extruded sample and the LS-3D printed sample where the drug exists as an amorphous solid dispersion. Furthermore, the role of polymer to inhibit precipitation and sustain the supersaturation conditions achieved by silicon dioxide-based formulations have been previously demonstrated [60]. Which explains the stability of the amorphous drug in the acidic medium before the pH shift and the stable ‘spring-parachute’ effect observed for the IND-inorganic carrier-Kollidon^®^ VA64 system both LS 3D printed and hot melt extrusion-based.

## 5. Conclusion

Bulk properties, especially the flow properties of pharmaceutical powders are extremely important for the smooth mass transfer of material during pharmaceutical manufacturing, drug content uniformity of the dosage forms, and the reproducibility of the process. These concepts also extend to novel manufacturing techniques of pharmaceutical dosage forms which include powder bed-based AM platforms such as powder bed fusion and binder jetting. In this study, we successfully demonstrated a novel granulation technique for developing powders for the LS 3D printing process. Further, it was observed that the processed powder not only exhibited excellent flow properties, and morphological properties but also demonstrated quicker dissolution behavior (>80% release in 5 minutes) attributed to the amorphous conversion of the crystalline drug (60% release in 2 hours). This conversion was depicted using polarized light microscopy where post-processing the samples did not exhibit any birefringence. These qualitative observations were further inspected using powder X-ray diffraction and modulated differential scanning calorimetric analysis. The changes in the intermolecular interactions post-processing were investigated using Fourier-transform infrared spectroscopy and Raman spectroscopy. The FT-IR and Raman spectra provided important information to understand the stability of the drug in acidic and the basic environment during the *in vitro* dissolution testing. Furthermore, the manufactured granules were successfully used for LS 3D printing tablets. The granulation process and the LS 3D printing process were found to be reproducible as all the quality control parameters for the granules and the manufactured tablets depicted a narrow standard deviation with insignificant variance. This technique can find applicability in manufacturing functional granules with enhanced performance for powder bed-based 3D printing techniques and expand their scope in pharmaceutical dosage form manufacturing.

## Funding information

The research work reported herein was supported by Maniruzzaman’s start-up funds at The University of Texas at Austin, and the Faculty Science and Technology Acquisition and Retention (STARs) Award. The authors and specifically Rishi Thakkar and Jiaxiang Zhang would also like to acknowledge the financial support from CoM3D Ltd., under an existing Master Sponsored Research Agreement (UTA19-000358) with UT Austin.

## Conflicts of Interest & Disclosures

The authors declare no conflict of interest. Rishi Thakkar, Jiaxiang Zhang, and Mohammed Maniruzzaman are co-investors on the related intellectual property.

## Notes

### Competing Interest Statement

The authors have declared no competing interest.

## References

1. Ngo, T. D., Kashani, A., Imbalzano, G., Nguyen, K. T. Q. & Hui, D. Additive manufacturing (3D printing): A review of materials, methods, applications and challenges. Composites Part B: Engineering 143, 172–196 (2018).

2. Ziaee, M. & Crane, N. B. Binder jetting: A review of process, materials, and methods. Additive Manufacturing 28, 781–801 (2019).

3. Fu, J., Yu, X. & Jin, Y. 3D printing of vaginal rings with personalized shapes for controlled release of progesterone. International Journal of Pharmaceutics 539, 75–82 (2018).

4. Lamichhane, S., Park, J.-B., Sohn, D. H. & Lee, S. Customized Novel Design of 3D Printed Pregabalin Tablets for Intra-Gastric Floating and Controlled Release Using Fused Deposition Modeling. Pharmaceutics 11, (2019).

5. Trenfield, S. J. et al. 3D printed drug products: Non-destructive dose verification using a rapid point-and-shoot approach. International Journal of Pharmaceutics 549, 283–292 (2018).

6. Goyanes, A., Det-Amornrat, U., Wang, J., Basit, A. W. & Gaisford, S. 3D scanning and 3D printing as innovative technologies for fabricating personalized topical drug delivery systems. J Control Release 234, 41–48 (2016).

7. Zhang, J., Thakkar, R., Zhang, Y. & Maniruzzaman, M. Structure-Function Correlation and Personalized 3D Printed Tablets using a Quality by Design (QbD) Approach. International Journal of Pharmaceutics 119945 (2020) doi:10.1016/j.ijpharm.2020.119945.

8. Jiang, J., Xu, X. & Stringer, J. Support Structures for Additive Manufacturing: A Review. Journal of Manufacturing and Materials Processing 2, 64 (2018).

9. Chai, X. et al. Fused Deposition Modeling (FDM) 3D Printed Tablets for Intragastric Floating Delivery of Domperidone. Sci Rep 7, 2829 (2017).

10. Yan, T.-T. et al. Semi-solid extrusion 3D printing ODFs: an individual drug delivery system for small scale pharmacy. Drug Development and Industrial Pharmacy 46, 531–538 (2020).

11. Davis, D. A., Thakkar, R., Su, Y., Williams, R. O. & Maniruzzaman, M. Selective Laser Sintering 3-Dimensional Printing as a Single Step Process to Prepare Amorphous Solid Dispersion Dosage Forms for Improved Solubility and Dissolution rate. Journal of Pharmaceutical Sciences S0022354920307413 (2020) doi:10.1016/j.xphs.2020.11.012.

12. Jacob, J., Caputo, K., Guillot, M., Sultzbaugh, K. J. & West, T. G. Rapidly dispersible dosage form of oxcarbazepine. (2016).

13. Thakkar, R. et al. Novel On-Demand 3-Dimensional (3-D) Printed Tablets Using Fill Density as an Effective Release-Controlling Tool. Polymers 12, 1872 (2020).

14. Dores, F. et al. Temperature and solvent facilitated extrusion based 3D printing for pharmaceuticals. European Journal of Pharmaceutical Sciences 152, 105430 (2020).

15. Fina, F., Goyanes, A., Gaisford, S. & Basit, A. W. Selective laser sintering (SLS) 3D printing of medicines. International Journal of Pharmaceutics 529, 285–293 (2017).

16. Chang, S.-Y. et al. Binder-Jet 3D Printing of Indomethacin-laden Pharmaceutical Dosage Forms. Journal of Pharmaceutical Sciences 109, 3054–3063 (2020).

17. Tan, Y., Zheng, J., Gao, W., Jiang, S. & Feng, Y. The Effect of Powder Flowability in the Selective Laser Sintering Process. in Proceedings of the 7th International Conference on Discrete Element Methods (eds. Li, X., Feng, Y. & Mustoe, G.) 629–636 (Springer, 2017). doi:10.1007/978-981-10-1926-5_65.

18. Muselík, J. et al. Influence of Process Parameters on Content Uniformity of a Low Dose Active Pharmaceutical Ingredient in a Tablet Formulation According to GMP. Acta Pharmaceutica 64, 355–367 (2014).

19. Kruth, J. P., Wang, X., Laoui, T. & Froyen, L. Lasers and materials in selective laser sintering. Assembly Automation 23, 357–371 (2003).

20. Salmoria, G. V., Klauss, P., Zepon, K. M. & Kanis, L. A. The effects of laser energy density and particle size in the selective laser sintering of polycaprolactone/progesterone specimens: morphology and drug release. Int J Adv Manuf Technol 66, 1113–1118 (2013).

21. Schmid, M., Amado, A. & Wegener, K. Polymer powders for selective laser sintering (SLS). in 160009 (2015). doi:10.1063/1.4918516.

22. Megarry, A. J., Swainson, S. M. E., Roberts, R. J. & Reynolds, G. K. A big data approach to pharmaceutical flow properties. International Journal of Pharmaceutics 555, 337–345 (2019).

23. Kashani Rahimi, S., Paul, S., Sun, C. C. & Zhang, F. The role of the screw profile on granular structure and mixing efficiency of a high-dose hydrophobic drug formulation during twin screw wet granulation. International Journal of Pharmaceutics 575, 118958 (2020).

24. Kittikunakorn, N., Sun, C. C. & Zhang, F. Effect of screw profile and processing conditions on physical transformation and chemical degradation of gabapentin during twin-screw melt granulation. European Journal of Pharmaceutical Sciences 131, 243–253 (2019).

25. Seem, T. C. et al. Twin screw granulation — A literature review. Powder Technology 276, 89–102 (2015).

26. Bardin, M., Knight, P. C. & Seville, J. P. K. On control of particle size distribution in granulation using high-shear mixers. Powder Technology 140, 169–175 (2004).

27. Jacob, J. et al. Rapid disperse dosage form. (2017).

28. Shah, V. P. & Amidon, G. L. G. L. Amidon, H. Lennernas, V.P. Shah, and J.R. Crison. A Theoretical Basis for a Biopharmaceutic Drug Classification: The Correlation of In Vitro Drug Product Dissolution and In Vivo Bioavailability, Pharm Res 12, 413–420, 1995—Backstory of BCS. AAPS J 16, 894–898 (2014).

29. Petruševska, M. et al. Biowaiver Monographs for Immediate Release Solid Oral Dosage Forms: Levetiracetam. Journal of Pharmaceutical Sciences 104, 2676–2687 (2015).

30. Crowley, M. M. et al. Pharmaceutical Applications of Hot-Melt Extrusion: Part I. Drug Development and Industrial Pharmacy 33, 909–926 (2007).

31. PubChem. Indomethacin. https://pubchem.ncbi.nlm.nih.gov/compound/3715 (2020).

32. Semjonov, K. et al. Interdependence of particle properties and bulk powder behavior of indomethacin in quench-cooled molten two-phase solid dispersions. International Journal of Pharmaceutics 541, 188–197 (2018).

33. Zeleňák, V. et al. Ordered cubic nanoporous silica support MCM-48 for delivery of poorly soluble drug indomethacin. Applied Surface Science 443, 525–534 (2018).

34. Tanabe, S. et al. Yellow coloration phenomena of incorporated indomethacin into folded sheet mesoporous materials. International Journal of Pharmaceutics 429, 38–45 (2012).

35. Taylor, M. K., Ginsburg, J., Hickey, A. J. & Gheyas, F. Composite method to quantify powder flow as a screening method in early tablet or capsule formulation development. AAPS PharmSciTech 1, 20–30 (2000).

36. Nováková, L., Matysová, L., Havlíková, L. & Solich, P. Development and validation of HPLC method for determination of indomethacin and its two degradation products in topical gel. Journal of Pharmaceutical and Biomedical Analysis 37, 899–905 (2005).

37. Ma, X. & Williams, R. O. Characterization of amorphous solid dispersions: An update. Journal of Drug Delivery Science and Technology 50, 113–124 (2019).

38. Otsuka, M., Kato, F. & Matsuda, Y. Comparative evaluation of the degree of indomethacin crystallinity by chemoinfometrical Fourier-transformed [corrected] near-infrared spectroscopy and conventional powder X-ray diffractometry. AAPS PharmSci 2, E9 (2000).

39. Davis, D. A., Miller, D. A. & Williams, R. O. Thermally conductive excipient expands kinetiSol^®^ processing capabilities. AAPS PharmSciTech **In press**, (2020).

40. Abiad, M. G., Carvajal, M. T. & Campanella, O. H. A Review on Methods and Theories to Describe the Glass Transition Phenomenon: Applications in Food and Pharmaceutical Products. Food Eng. Rev. 1, 105–132 (2009).

41. Shimada, Y. et al. Decarboxylation of indomethacin induced by heat treatment. International Journal of Pharmaceutics 545, 51–56 (2018).

42. Garbacz, P. & Wesolowski, M. DSC, FTIR and Raman Spectroscopy Coupled with Multivariate Analysis in a Study of Co-Crystals of Pharmaceutical Interest. Molecules 23, 2136 (2018).

43. Ewing, A. V., Clarke, G. S. & Kazarian, S. G. Stability of indomethacin with relevance to the release from amorphous solid dispersions studied with ATR-FTIR spectroscopic imaging. European Journal of Pharmaceutical Sciences 60, 64–71 (2014).

44. Taylor, L. S. & Zografi, G. Spectroscopic Characterization of Interactions Between PVP and Indomethacin in Amorphous Molecular Dispersions. Pharm Res 14, 1691–1698 (1997).

45. Hédoux, A., Paccou, L., Guinet, Y., Willart, J.-F. & Descamps, M. Using the low-frequency Raman spectroscopy to analyze the crystallization of amorphous indomethacin. European Journal of Pharmaceutical Sciences 38, 156–164 (2009).

46. O’Brien, M., McCauley, J. & Cohen, E. Indomethacin. in Analytical Profiles of Drug Substances (ed. Florey, K.) vol. 13 211–238 (Academic Press, 1984).

47. Dubbini, A. et al. Influence of pH and method of crystallization on the solid physical form of indomethacin. International Journal of Pharmaceutics 473, 536–544 (2014).

48. Jermain, S. V. et al. In Vitro and In Vivo Behaviors of KinetiSol and Spray-Dried Amorphous Solid Dispersions of a Weakly Basic Drug and Ionic Polymer†. Molecular Pharmaceutics (2020) doi:10.1021/acs.molpharmaceut.0c00108.

49. Skrdla, P. J., Floyd, P. D. & Dell’Orco, P. C. Modeling Recrystallization Kinetics Following the Dissolution of Amorphous Drugs. Mol. Pharmaceutics 17, 219–228 (2020).

50. Huang, S., Mao, C., Williams, R. O. & Yang, C.-Y. Solubility Advantage (and Disadvantage) of Pharmaceutical Amorphous Solid Dispersions. J Pharm Sci 105, 3549–3561 (2016).

51. Maniruzzaman, M. et al. Drug–polymer intermolecular interactions in hot-melt extruded solid dispersions. International Journal of Pharmaceutics 443, 199–208 (2013).

52. Nie, H. et al. Investigating the Interaction Pattern and Structural Elements of a Drug–Polymer Complex at the Molecular Level. Mol. Pharmaceutics 12, 2459–2468 (2015).

53. Viidik, L. et al. Preparation and characterization of hot-melt extruded polycaprolactone-based filaments intended for 3D-printing of tablets. European Journal of Pharmaceutical Sciences 105619 (2020) doi:10.1016/j.ejps.2020.105619.

54. Xu, P. et al. Development of a quantitative method to evaluate the printability of filaments for fused deposition modeling 3D printing. International Journal of Pharmaceutics 588, 119760 (2020).

55. Li, Y., Rantanen, J., Yang, M. & Bohr, A. Molecular structure and impact of amorphization strategies on intrinsic dissolution of spray dried indomethacin. European Journal of Pharmaceutical Sciences 129, 1–9 (2019).

56. Xi, Z. et al. Evaluation of the Solid Dispersion System Engineered from Mesoporous Silica and Polymers for the Poorly Water Soluble Drug Indomethacin: In Vitro and In Vivo. Pharmaceutics 12, 144 (2020).

57. Majumder, M., Rajabnezhad, S., Nokhodchi, A. & Maniruzzaman, M. Chemico-calorimetric analysis of amorphous granules manufactured via continuous granulation process. Drug Deliv Transl Res 8, 1658–1669 (2018).

58. Mohapatra, S., Samanta, S., Kothari, K., Mistry, P. & Suryanarayanan, R. Effect of Polymer Molecular Weight on the Crystallization Behavior of Indomethacin Amorphous Solid Dispersions. Crystal Growth & Design 17, 3142–3150 (2017).

59. Kosaka, M., Higashi, K., Nishimura, M., Ueda, K. & Moribe, K. Clarification of the Dissolution Mechanism of an Indomethacin/Saccharin/Polyvinylpyrrolidone Ternary Solid Dispersion by NMR Spectroscopy. Journal of Pharmaceutical Sciences 109, 3617–3624 (2020).

60. Hanada, M., Jermain, S. V. & Williams, R. O. Enhanced Dissolution of a Porous Carrier–Containing Ternary Amorphous Solid Dispersion System Prepared by a Hot Melt Method. Journal of Pharmaceutical Sciences 107, 362–371 (2018).

